# Multidimensional host-associated diversification in natural *Festuca–Epichloë festucae* symbioses across the Iberian Peninsula

**DOI:** 10.64898/2026.08.22.746409

**Authors:** Alba Sotomayor-Alge, Padmaja Nagabhyru, Beatriz R. Vázquez de Aldana, Luis A. Inda, Íñigo Zabalgogeazcoa, Christopher L. Schardl, Pilar Catalán

## Abstract

*Epichloë* fungal endophytes form widespread symbioses with temperate grasses, yet the extent to which diversity within endophyte species is shaped by host association remains poorly understood. Here, we characterized naturally occurring *Festuca–Epichloë* symbioses across diverse Iberian ecosystems using an integrative framework combining ecological, cytogenetic, phenotypic, molecular and chemical analyses. Novel associations of *Epichloë festucae* with *Festuca trichophylla*, *F. lambinonii* and *F. yvesii* were documented, together with substantial variation in infection incidence and mating-type composition among host-associated populations. Morphological traits, vegetative growth and alkaloid profiles differentiated strains according to host identity. Furthermore, multilocus phylogenetic analyses assigned all fine-leaved *Festuca* host isolates to *Epichloë festucae*, but identified a recurrent host-associated genetic structure, along with a deeper evolutionary signal, that largely corresponds to the host phylogeny. By contrast, genome size estimates varied little among *Epichloë festucae* strains, with all isolates exhibiting haploid genomes. Alkaloid content across the four major classes of *Epichloë* compounds (pyrrolopyrazines, 1-aminopyrrolizidines, ergot alkaloids and indole-diterpenes) showed only partial concordance with the presence of biosynthetic genes, indicating that functional outcomes are influenced by regulatory and environmental factors beyond biosynthetic gene presence. Chemotypic profiles clearly differentiated *Epichloë festucae* from *E. coenophiala* while demonstrating considerable functional diversity among *E. festucae* strains. Collectively, these complementary datasets reveal two interconnected signatures of diversification: pervasive host-associated differentiation across multiple biological dimensions and a deeper historical signal retained in phylogenetic relationships. These findings provide a foundation for future genomic, evolutionary and systematic studies to determine whether these lineages represent ongoing fungal divergence and speciation.

**Significance statement:** Understanding how diversity is generated and maintained within plant-associated symbionts remains a fundamental challenge in biology. By integrating ecological, phylogenetic, phenotypic and chemical evidence across natural *Festuca–Epichloë* symbioses, including newly documented host associations, we reveal two complementary signatures of host-associated diversification within a widespread fungal endophyte. Beyond expanding the known diversity of these symbioses, our findings highlight natural *Festuca*–*Epichloë* associations as a valuable framework for exploring diversification in plant–microbe interactions.

## INTRODUCTION

*Epichloë* species (*Clavicipitaceae*, *Ascomycota*; Leuchtmann et al., 2014) form biotrophic associations with cool-season *Pooideae* grasses (Schardl, 1996; Clay & Schardl, 2002; Moon et al., 2004; Gentile et al., 2005; Leuchtmann and Schardl, 2005). These symbioses involve systemic colonization of aerial vegetative and reproductive host tissues, where the endophyte may engage in asexual, sexual, or mixed reproductive strategies. Vertical transmission occurs innocuously through host seeds and horizontal transmission takes place via stromata formation on host inflorescences (Leuchtmann et al., 2014; Tadych et al., 2014). While many *Epichloë* species can only reproduce asexually, others retain the capacity for sexual reproduction, reflecting a diversity of life-history strategies within the genus. These differences in transmission mode and reproductive strategy influence patterns of gene flow and may contribute to diversification within *Epichloë* lineages (Schardl et al., 2004; Saikkonen et al., 2016).

*Epichloë*-grass symbioses span a continuum from mutualism to antagonism but are most commonly described by the scientific community as ’defensive mutualisms’ (Saikkonen et al., 2013), owing to the endophyte biochemical activities, which have been shown to enhance resistance against biotic stresses through the production of secondary metabolites, hormonal crosstalk, and complex physiological responses (e.g., Xia et al., 2018; Bastías and Gundel, 2023). Specifically, four major families of alkaloids have been identified in *Epichloë* (Schardl et al., 2013). Among them, indole-diterpenes (e.g., lolitrem B) and ergot alkaloids (e.g., ergovaline) are known for their potent toxicity to vertebrate herbivores, whereas 1-aminopyrrolizidines (e.g., lolines) and pyrrolopyrazines (e.g., peramine) act as well- established bioinsecticides or feeding deterrents, and some non-tremorgenic indole-diterpenes (e.g., terpendole E) are associated with anti-insect activity and reduced vertebrate toxicity (Saikkonen et al., 2013; Bastías et al., 2017; Vassiliadis et al., 2023). Each alkaloid family targets distinct physiological systems in herbivores, and its biosynthesis is tightly regulated and variable (Zhang et al., 2009; Chujo and Scott, 2014 Panaccione et al., 2014; Guerre, 2016; Caradus and Johnson, 2020). Owing to their pronounced effects on livestock health and crop protection and improvement, these compounds have garnered significant attention in agriculturally important grass species such as *Lolium perenne* L. and *Festuca arundinacea* L. over recent decades (e.g., Bacon et al., 1977; Clay, 1988; Schardl et al., 2004; Young et al., 2013; Johnson et al., 2013; Hume and Sewell, 2014; Ferguson et al., 2021; Card et al., 2024). However, few studies have systematically explored the extent of intraspecific genetic, phenotypic and functional variation within *Epichloë* species inhabiting naturally distributed grass hosts.

*Festuca* L. is one of the most ecologically and economically important worldwide distributed grass genera within the subtribe Loliinae (Catalán, 2006). It comprises more than 600 species (Catalán, 2006; Inda et al., 2008; Moreno-Aguilar et al., 2026) and includes many relevant forage, pasture, and lawn grasses (Kopecký and Studer, 2014). Recent phylogenetic studies on *Festuca* and closely related genera (e.g., Inda et al., 2008; Minaya et al., 2017; Moreno-Aguilar et al., 2020; Moreno-Aguilar et al., 2022; Moreno-Aguilar et al., 2026) support the divergence of two major fine-leaved (FL) and broad-leaved (BL) Loliinae clades. Specifically, *Epichloë festucae* is known to inhabit a wide range of fescues belonging to the FL Loliinae clade (e.g., Leuchtmann et al., 1994; Naffaa et al., 1998; Moon et al., 2004; Zabalgogeazcoa et al., 2006; Gibert and Hazard, 2011; Gundel et al., 2014; Vázquez de Aldana et al., 2015; Dirihan et al., 2016; von Cräutlein et al., 2021), and a few species from the BL Loliinae (Bush et al., 1997; Cagnano et al., 2019) and the intermediate Loliinae clades (Niones and Takemoto, 2014). This broad host distribution provides a valuable framework for exploring patterns of variation within this widespread *Epichloë* species across diverse host backgrounds. Despite this potential, the extent of genetic, phenotypic and functional variation within *E. festucae* across different *Festuca* hosts remains poorly characterized.

Understanding patterns of diversity within *Epichloë*–grass symbioses require consideration of both symbiotic partners. Host diversity can be characterized using morphological, cytogenetic and molecular approaches (e.g., Dirihan et al., 2016; Moreno-Aguilar et al., 2022b; Sotomayor-Alge et al., 2025; Moreno-Aguilar et al., 2026), whereas endophyte characterization additionally benefits from reproductive, phylogenetic and alkaloid profiling, which together provide complementary insights into their ecological and functional diversity (e.g., Gentile et al., 2005; Zabalgogeazcoa et al., 2006; McCargo et al., 2014; Vikuk et al., 2019; Du et al., 2024). Together, these complementary approaches enable a more comprehensive assessment of how genetic, phenotypic and functional diversity are distributed at species level and whether such variation is associated with different host backgrounds.

Given the broad host range of *Epichloë festucae* and the ecological and evolutionary diversity of its *Festuca* hosts, we hypothesized that endophyte strains associated with different host species may exhibit host-associated genetic, phenotypic and functional variation. To test this hypothesis, we investigated six naturally occurring *Festuca–Epichloë* holobionts distributed across contrasting environments of the Iberian Peninsula, including the first characterization of *Epichloë* associations in *Festuca trichophylla*, *F. lambinonii* and *F. yvesii*, and a broader assessment of this symbiosis in *F. rothmaleri*, *F. nigrescens* and *F. rubra* subsp. *pruinosa*. Using an integrative framework combining morphological, molecular, cytogenetic and biochemical approaches, we characterized variation within *Epichloë festucae* and evaluated whether patterns of genetic, phenotypic and functional diversity are associated with host species.

## RESULTS

### Endophyte incidence, and reproductive and cytogenetic variation among *Festuca*– *Epichloë* holobionts

*Epichloë* detection was consistent between aniline blue staining and direct isolation on Potato Dextrose Agar (PDA) medium. Endophyte incidence varied markedly among host species, ranging from complete absence in *Festuca ampla*, *F. elegans*, and *F. paniculata* (0%) and very low incidence in *F. trichophylla* and *F. lambinonii* (<5%), to relatively high incidence in *F. yvesii* (60%) and *F. rubra* subsp. *pruinosa* (65%). *F. nigrescens* and *F. arundinacea* subsp. *arundinacea* exhibited intermediate incidence values of approximately 50% (Table 1). Hereafter, the fine-leaved holobionts are referred to as Flamb (*F. lambinonii*–*E. festucae*), Fyves (*F. yvesii*–*E. festucae*), Ftrich (*F. trichophylla*–*E. festucae*), Fnigr (*F. nigrescens*–*E. festucae*) and Fprui (*F. rubra* subsp. *pruinosa* – *E. festucae*). Additionally, the previously characterized *F. rothmaleri*–*E. festucae* holobiont (Sotomayor-Alge et al., 2025) is referred to as Froth, and the *F. arundinacea* subsp. *arundinacea*–*E. coenophiala* holobiont as Farun.

**Table 1.** Fine-leaved and broad-leaved *Festuca-Epichloë* holobionts included in the comparative study of host-endophyte interactions. Summary of sampling campaigns conducted in northeastern and northwestern Spain. Sample information (host species and population IDs), collection metadata (province of origin, coordinates, sampling dates and reference), environmental characteristics (site ecology and altitude in meters above sea level), and endophyte presence (*Epichloë* incidence (E^+^) and mating type composition of the endophyte populations). When both mating types were detected, it was among different individuals of the same population.

| Sample identification |  | Collection metadata |  |  |  |  | Environmental characteristics |  | Endophyte presence |  |
| --- | --- | --- | --- | --- | --- | --- | --- | --- | --- | --- |
| Host Species | Pop. IDs | Province | Latitude | Longitude | Date | Reference | Ecology | Altitude (m) | Incidence (%) | Mating type |
| *Festuca lambinonii* | UZ_13.22<br>UZ_3.23 | Huesca | 42°18'42.9"N | 0°22'55.4"W | June 2022<br>June 2023 | This study | Limestone and clay soils. Oak grasslands, old field margins with *Rosa* gr. *canina* | 1211 | 4.4%<br>(1/23) | MTA |
| *Festuca yvesii* | UZ_20.22<br>UZ_7.23 | Huesca | 42°19'57.5"N | 0°07'27.6"W | June 2022<br>June 2023 | This study | Limestone soils. Riverbanks with *Salix atrocinerea* and *Crataegus monogyna* | 1123 | 60%<br>(9/15) | MTA,<br>MTB |
| *Festuca trichophylla* | UZ_4.22<br>UZ_1.23 | Huesca | 41°48'55.4"N | 0°30'35.0"W | May 2022,<br>May 2023 | This study | Limestone and gypsum soils. Woods of *Acer monspessulanus* with *Rosa* gr. *canina* and *Prunus spinosa* | 596 | 4.2 %<br>(1/24) | MTB |
| *Festuca rothmaleri* | UZ_35.22<br>UZ_9.23<br>UZ_10.23<br>UZ_11.23<br>UZ_12.23 | Salamanca | 40°20'19.3"N | 5°46'00.9"W | June 2022<br>June 2023 | Sotomayor-Alge et al. (2025) | Granite, acidophilous and sandy soils. Riverbank with oak forest. | 1254 | 84%<br>(21/25) | MTA,<br>MTB |
| *Festuca nigrescens* | UZ_42.22<br>UZ_44.22<br>UZ_27.23 | Huesca | 42°45'09.8"N | 0°18'36.5"W | July 2022<br>July 2023 | This study | Mixed calcareous and siliceous soils. Mountain riparian and lacustrine ecosystem with hygrophilous herbaceous communities and low shrub vegetation | 1655 | 47.6%<br>(10/21) | MTA,<br>MTB |
| *Festuca rubra*<br>subsp. *pruinosa* | UZ_55.22<br>UZ_18.23 | La Coruña | 43°23'44.8"N | 8°20'56.3"W | June 2022<br>June 2023 | This study | Acidic, shallow soils exposed to wind and salt spray. Maritime heathland and grassland communities. | 36 | 50%<br>(7/14) | MTA |
|  | UZ_56.22<br>UZ_17.23 | La Coruña | 43°22'13.2"N | 8°27'19.4"W | June 2022<br>June 2023 | This study | Acidic, shallow soils exposed to wind and salt spray. Maritime heathland and grassland communities. | 15 | 80%<br>(8/10) | MTA |
| *Festuca arundinacea* subsp. *arundinacea* | UZ_11.22,<br>UZ_12.22<br>UZ_2.23 | Huesca | 42°18'42.9"N | 0°22'56.6"W | June 2022<br>June 2023 | This study | Limestone and clay soils. Stream banks with *Populus tremula*, *Euonymus europaeus*, *Acer campestre*, *Crataegus monogyna* and *Buxus sempervirens* | 1211 | 44.4%<br>(8/18) | MTA |
| *Festuca ampla* | UZ_41.22 | Salamanca | 40°29'12.7"N | 5°31'14.2"W | June 2022 | This study | Granite and sandy soils. Riverbank with herbaceous communities. | 959 | 0%<br>(0/15) | - |
| *Festuca elegans* | UZ_27.22 | Salamanca | 40°20'18.7"N | 5°46'01.3"W | June 2022 | This study | Granite, acidophilous and sandy soils. Oak forest. | 1254 | 0%<br>(0/10) | - |
| *Festuca paniculata* | UZ_29.22 | Salamanca | 40°20'17.5"N | 5°45'59.2"W | June 2022 | This study | Granite, acidophilous and sandy soils. Oak forest. | 1254 | 0%<br>(0/12) | - |

PCR screening of mating-type idiomorphs revealed contrasting reproductive patterns among holobionts. A single mating type was detected in Flamb (*MTA*), Ftrich (*MTB*), Fprui (*MTA*), and Farun (*MTA*), whereas both mating types were detected in different individuals of Fyves and Fnigr populations (Table 1). Stromata were occasionally observed in *F. nigrescens*, indicating that the coexistence of both mating types does not necessarily imply expression of the sexual stage under the conditions observed (White, 1988).

Flow cytometry analyses revealed substantial variation in host genome size among *Festuca* species (Table 2). Values for the three species from *Festuca* sect. *Aulaxyper* (*F. trichophylla*, *F. nigrescens* and *F. rubra* subsp. *pruinosa*) are consistent with hexaploid cytotypes (12.10– 12.74 pg/2C; Šmarda et al., 2008; Garnatje et al., 2023). In *Festuca* sect. *Festuca*, the value for *F. yvesii* (13.18 ± 0.35 pg/2C) falls within the range of hexaploid cytotypes, whereas that of *F. lambinonii*, with genome size reported here for the first time (9.19 pg/2C), is consistent with a tetraploid cytotype (Garnatje et al., 2023). Likewise, in broad-leaved *Festuca* subgen. *Schedonorus* the value for *F. arundinacea* subsp. *arundinacea* (16.17 pg/2C; *F.* subgen. *Schedonorus*) corresponds to an hexaploid cytotype (Šmarda et al., 2008) with larger genome size and distinct genomic composition relative to fine-leaved fescues (Moreno-Aguilar et al., 2022; Moreno-Aguilar et al., 2026). Coefficients of variation remained below 3% across all samples, except in *F. trichophylla*, where slightly higher values were observed. By contrast, genome size varied little among *Epichloë* isolates from fine-leaved hosts, ranging from 0.045 ± 0.004 pg/1C in Fprui to 0.050 ± 0.001 pg/1C in Ftrich, consistent with haploid genomes (Table 2). In contrast, *E. coenophiala* (Farun) displayed a markedly larger genome size (0.111 ± 0.007 pg/1C), approximately three times larger than those of fine-leaved-associated *Epichloë* strains and consistent with its heteroploid origin (Tsai et al., 1994). This provides the first flow cytometry-based evidence supporting the heteroploid nature previously reported for *E. coenophiala*.

**Table 2.** Genome size estimates obtained by flow cytometry for *Festuca* host plants and their associated *Epichloë* endophytes. Holobionts analyzed belong to fine-leaved *Festuca* sect. *Festuca* [*F*. *lambinonii* (Flamb), *F. yvesii* (Fyves)], and *Festuca* sect. *Aulaxyper* [*F. trichophylla* (Ftrich), *F*. *nigrescens* (Fnigr), *F*. *rubra* subsp. *pruinosa* (Fprui)], and broad-leaved *Festuca* subgen. *Schedonorus* [*F*. *arundinacea* subsp. *arundinacea* (Farun)]. N indicates the number of genetically independent host individuals or fungal isolates analyzed in this study. For each individual or isolate, two technical replicates were performed, resulting in a total of 2N cytometric measurements used to estimate genome size. In those cases where N = 1, six technical replicates were obtained from the available individual following standard flow cytometry procedures. Values represent species mean genome sizes ± SD, expressed as pg/2C for host species and pg/1C for endophytes. The coefficient of variation (CV) shown corresponds to the average CV calculated for each case studied. Ploidy shows the inferred ploidy levels for *Festuca* host plants (4x, tetraploid; 6x, hexaploid) and for *Epichloë* isolates (1x, haploid; 3x, triheteroploid) based on the respective genome size estimations and previous studies (Garnatje et al., 2023; Sotomayor-Alge et al., 2025).

| Holobiont | Plant host |  |  |  |  | Fungal endophyte |  |  |  |  |
| --- | --- | --- | --- | --- | --- | --- | --- | --- | --- | --- |
|  | Species | N | pg/2C | CV | Ploidy | Species | N | pg/1C | CV | Ploidy |
| Flamb | *F. lambinonii* | 5 | 9.19<br>± 0.07 | 2.78 | 4x | *E. festucae* | 1 | 0.048<br>± 0.002 | 6.55 | 1x |
| Fyves | *F. yvesii* | 5 | 13.18<br>± 0.35 | 2.46 | 6x | *E. festucae* | 3 | 0.046<br>± 0.001 | 6.57 | 1x |
| Ftrich | *F. trichophylla* | 3 | 12.10<br>± 0.28 | 3.21 | 6x | *E. festucae* | 1 | 0.050<br>± 0.001 | 6.33 | 1x |
| Fnigr | *F. nigrescens* | 5 | 12.69<br>± 0.27 | 2.69 | 6x | *E. festucae* | 3 | 0.049<br>± 0.002 | 6.22 | 1x |
| Fprui | *F. rubra* subsp.<br>*pruinosa* | 5 | 12.74<br>± 0.21 | 2.25 | 6x | *E. festucae* | 3 | 0.045<br>± 0.004 | 6.73 | 1x |
| Farun | *F. arundinacea*<br>subsp. *arundinacea* | 5 | 16.17<br>± 0.12 | 2.53 | 6x | *E. coenophiala* | 3 | 0.111<br>± 0.007 | 4.74 | 3x |

### Morphological and growth diversity among *Epichloë* strains

Comparative analyses revealed substantial morphological variation among *Epichloë festucae* strains associated with different *Festuca* hosts. Conidial length and area had the strongest differentiation among strains, whereas conidial width displayed lower variability and substantial overlap (Figure 1A; Table S1). Ftrich exhibited the smallest conidia (4.0 ± 0.4 µm length; 6.4 ± 0.9 µm² area), whereas Fprui had the largest (5.0 ± 0.5 µm; 8.6 ± 1.1 µm²), with the remaining strains displaying intermediate values. By contrast, conidiogenous cell length followed a different pattern, with the lowest values observed in Fnigr (10.7 ± 2.5 µm) and the highest in Ftrich (16.0 ± 3.6 µm; Figure 1A). *E. coenophiala* (Farun) consistently had larger values for length-related traits than all *E. festucae* strains, whereas width-related traits showed substantial overlap (Table S1). To focus on intraspecific variation within *E. festucae*, *E. coenophiala* was excluded from subsequent statistical analyses.

**Figure 1.**
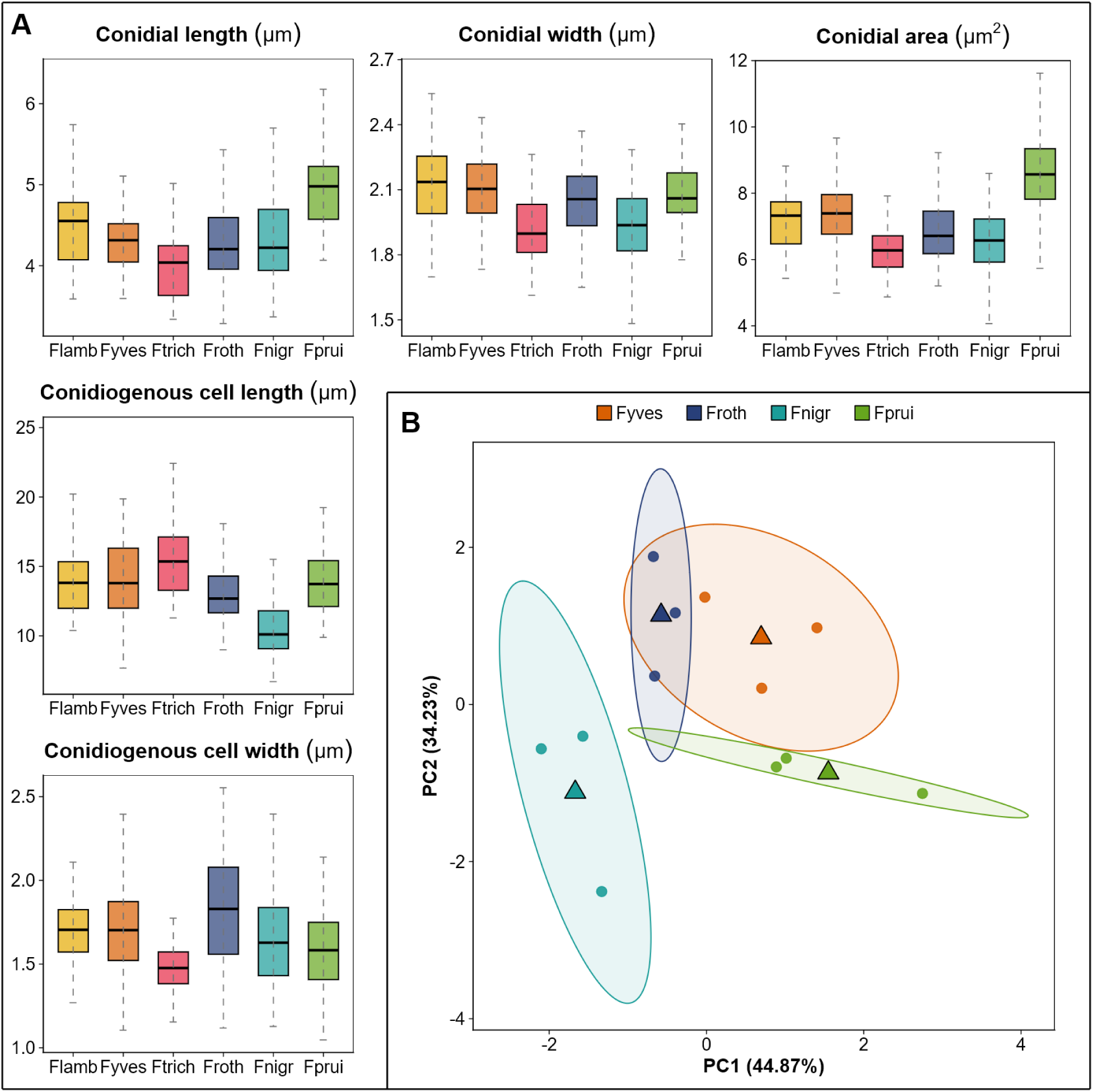
**(A)** Boxplots illustrating variation in morphological traits (conidial length, width and area, as well as conidiogenous cell length and width) among *Epichloë festucae* strains associated with different fine-leaved *Festuca* host species: *F.* sect. *Festuca*: *F. lambinonii* (Flamb; n = 30) and *F. yvesii* (Fyves n = 90); *F.* sect. *Aulaxyper*: *F. trichophylla* (Ftrich; n = 30), *F. rothmaleri* (Froth; n = 90), *F. nigrescens* (Fnigr; n = 90) and *F. rubra* subsp. *pruinosa* (Fprui; n = 90). Boxes represent the interquartile range (IQR), central lines indicate medians, and whiskers extend to the range of observed values (see also Table S1). **(B)** Principal component analysis (PCA) based only on independent morphological traits (conidial and conidiogenous cell dimensions, including length and width) of *E. festucae* strains from *Festuca* host species with three fungal isolates: *F. yvesii* (Fyves), *F. rothmaleri* (Froth), *F. nigrescens* (Fnigr) and *F. rubra* subsp. *pruinosa* (Fprui). Points represent individual fungal isolates colored according to holobiont identity, as shown in the chart. Ellipses indicate the dispersion of each group (95%), and triangles represent group centroids (see also Table S5).

Linear mixed-effects models broadly supported these descriptive patterns (Tables S2–S3). Host species had a significant effect on most morphological traits, including conidial width and area, and conidiogenous cell length and width, whereas the effect on conidial length was only marginal (*F*3,8 = 3.98, *p* = 0.053). Pairwise comparisons indicated that the strongest differences involved the *Epichloë festucae* strains associated with Fprui and Fnigr holobionts and, although width-related traits had lower variation in the descriptive analyses, they captured consistent differences among *Epichloë* strains (Table S2). Variance partitioning further indicated that the magnitude of inter-strain differentiation varied among traits, with conidial area providing the strongest support for differentiation among strains (*R*²m = 0.35), followed by conidiogenous cell length (*R*²m = 0.27) and conidial width (*R*²m = 0.12). Nevertheless, most of the total variation was attributable to residual variance, indicating substantial intra-isolate variability (Table S3).

Complementary multivariate analyses (PERMANOVA and PCA) further supported host-associated structuring of morphological variation in the asexual structures of *Epichloë festucae* strains (Figure 1B; Tables S4–S5). Global PERMANOVA detected significant differences among species at the isolate level (pseudo-*F*3,8 = 5.15, R² = 0.659, *p* < 0.001; Table S4A). Tests for homogeneity of multivariate dispersion were not significant, reflecting shifts in holobiont centroids rather than dispersion effects. Moreover, the proportion of explained variance increased from spore-level measurements (R² = 0.18) to isolate means (R² = 0.66), suggesting that within-isolate variability partially masks inter-holobiont differences at finer hierarchical levels. Pairwise PERMANOVA comparisons between holobionts yielded moderate to high effect sizes (R² = 0.41–0.68), with the strongest differentiation between Fprui and Froth and the weakest between Froth and Fyves (Table S4B). However, none of the pairwise comparisons remained significant after permutation testing, likely due to the limited number of isolates per species (n = 3). The PCA based on isolate means further supported these patterns, revealing partially differentiated morphospaces in which holobionts differed significantly along both PC1 (*F*₃,₈ = 13.60, *p* = 0.002) and PC2 (*F*₃,₈ = 7.35, *p* = 0.011) axes (Figure 1B; Table S5). Consistent with PERMANOVA results, Fnigr was clearly separated from the remaining holobionts along PC1, whereas Fprui occupied the opposite region of the morphospace and Froth and Fyves largely overlapped (Figure 1B). The first two principal components explained 79.1% of the total variance. PC1 was mainly associated with size-related traits, showing high positive loadings for conidiogenous cell length (0.793), conidial width (0.771) and conidial length (0.634), and PC2 with conidiogenous cell width (0.793) and conidial length (−0.584), reflecting variation in shape and proportional traits (Table S5). Inclusion of Farun increased the variance explained by PC1 and clearly separated *E. coenophiala* from all *E. festucae* strains (Figure S1).

Noticeable differences in macroscopic colony morphology and radial growth trajectories were observed among *Epichloë* strains associated with different host species during 3 weeks of cultivation on PDA (Figure 2). Growth rate analyses based on colony diameter measurements revealed significant differences among *Epichloë festucae* strains associated with the four fine-leaved *Festuca* host species (pseudo-*F*3,8 = 6.61, *p* = 0.015; Table S6A). Fnigr isolates exhibited the highest growth rates (1.98 ± 0.2 mm/day), whereas those from Fprui showed the lowest values (0.9 ± 0.09 mm/day; Figure 2A–B). Pairwise comparisons confirmed these patterns (Table S6A). Although Flamb was excluded from formal statistical analyses because only one isolate was available, its observed growth rate was among the highest recorded (Figure 2). Inclusion of Farun accentuated these differences (pseudo-*F*₄,₁₀ = 17.64, *p* < 0.001; Table S6B), with *Epichloë coenophiala* isolates recurrently displaying markedly lower growth rates than all *E. festucae* strains. Macroscopic colony morphology also differed among holobionts, particularly in colony density, texture, and marginal growth patterns, which became progressively more evident after the second week of cultivation (Figure 2C).

**Figure 2.**
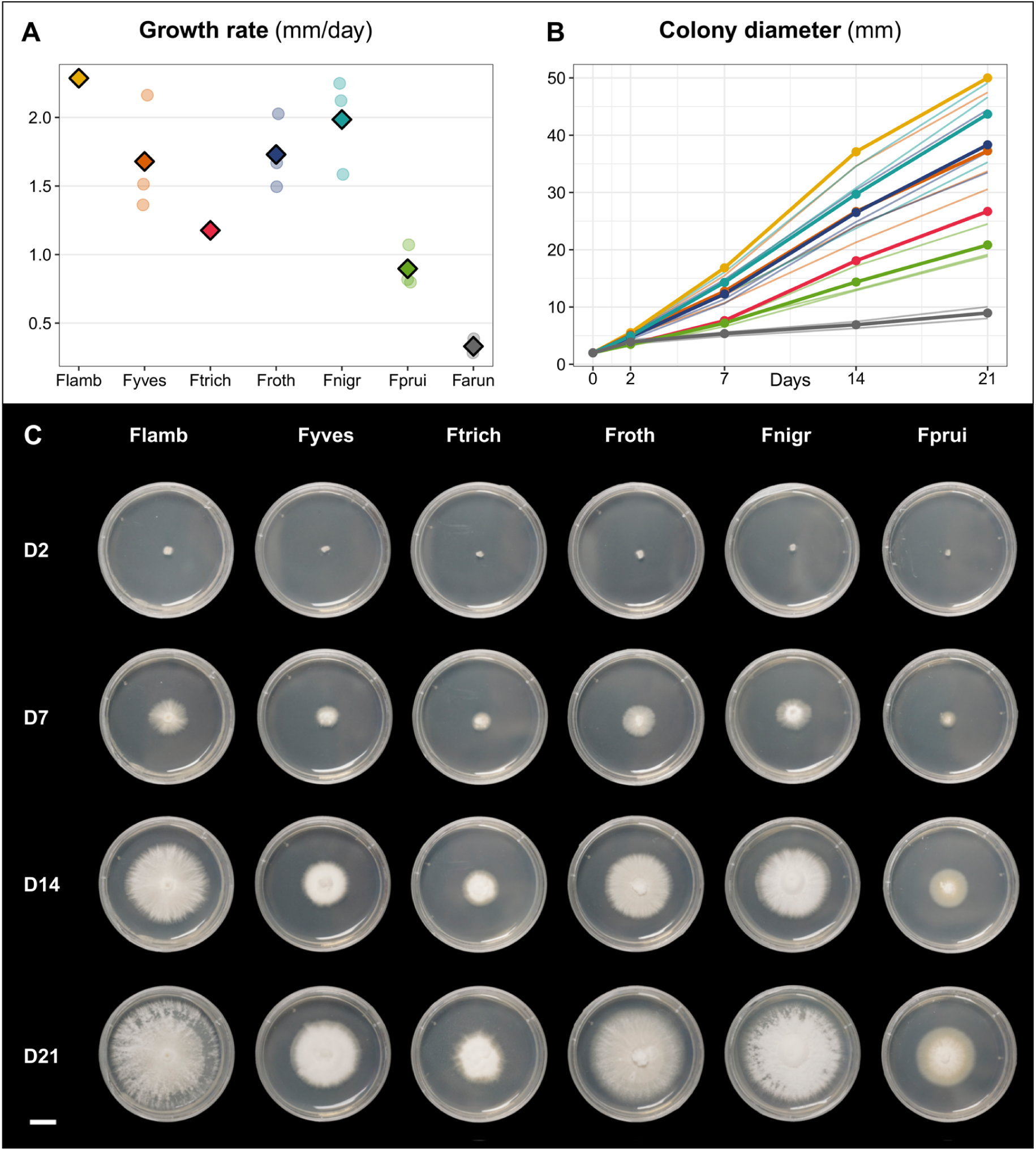
Growth characteristics of *Epichloë festucae* isolates cultured *in vitro*, isolated from fine-leaved *Festuca* host species: *F. lambinonii* (Flamb), *F. yvesii* (Fyves), *F. trichophylla* (Ftrich), *F. rothmaleri* (Froth), *F. nigrescens* (Fnigr) and *F. rubra* subsp. *pruinosa* (Fprui). **(A)** Culture growth rate for each isolate, with mean values (diamonds) highlighted for each holobiont. **(B)** Colony diameter over time (0, 2, 7, 14 and 21 days). Points represent individual measurements and lines track colony expansion dynamics. Colors associated with each *E. festucae* strain correspond to those of panel A. The heteroploid species *E. coenophiala* (*F. arundinacea* subsp. *arundinacea*; Farun) is also included in panels A and B. **(C)** Temporal reconstruction of culture growth of *E. festucae* strains on 5.5 cm PDA plates after 2, 7, 14 and 21 days. Isolates photographed are: UZ_13.22(5)-Flamb, UZ_7.23(11)-Fyves, UZ_4.22(1)-Ftrich, UZ_9.23(4)-Froth, UZ_44.22(4)-Fnigr, and UZ_55.22(4)-Fprui. For each *E. festucae* strain, the average growth rate is provided, calculated as mm/day ± SD. Scale bar: 1 cm.

### Phylogenetic relationships among *Epichloë* strains

Phylogenetic reconstructions based on five nuclear loci (*actG*, *CalM*, rDNA ITS region, *tefA*, *tubB*) using both multispecies coalescent (MSC; Figure 3;Tables S7-S8) and independent and concatenated multilocus (ML; Figures S2-S3; Table S8) approaches resolved strongly supported and broadly congruent topologies and consistently placed all newly sequenced isolates from fine-leaved *Festuca* holobionts within the *Epichloë festucae* clade (MSC: 0.99 PP and q1 = 1 in the MSC tree; ML: 100 BS and 71.1% SCFL). Although both approaches detected highly congruent topologies, they differed in the phylogenetic signal recovered within the *E. festucae* clade. The MSC reconstruction showed closer correspondence with the phylogenetic relationships previously described among their fine-leaved *Festuca* hosts, particularly the nesting of Flamb and Fyves holobionts’ strains within a subclade of *E. festucae* “*F.* sect. *Festuca”* and the separation of the Froth lineage from the remaining Fnigr, Ftrich and Fprui lineages of the *E. festucae* “*F*. sect. *Aulaxyper”* subclade, in parallel with those of their respective hosts (Inda et al., 2008; Minaya et al., 2017; Moreno-Aguilar et al., 2026). However, support values for several of these internal relationships remained moderate (PP < 0.87; Figure 3). In contrast, independent and concatenated ML phylogenies revealed different relationships (Figures S2-S3), with the latter showing higher branch support values for all pairs of isolates associated with the same holobiont (BS > 61), but with a close, though different, phylogenetic position for Ftrich and a non-monophyletic resolution for the Flamb and Fyves lineages of the *E. festucae* “*F.* sect. *Festuca”* group (Figure S3).

**Figure 3.**
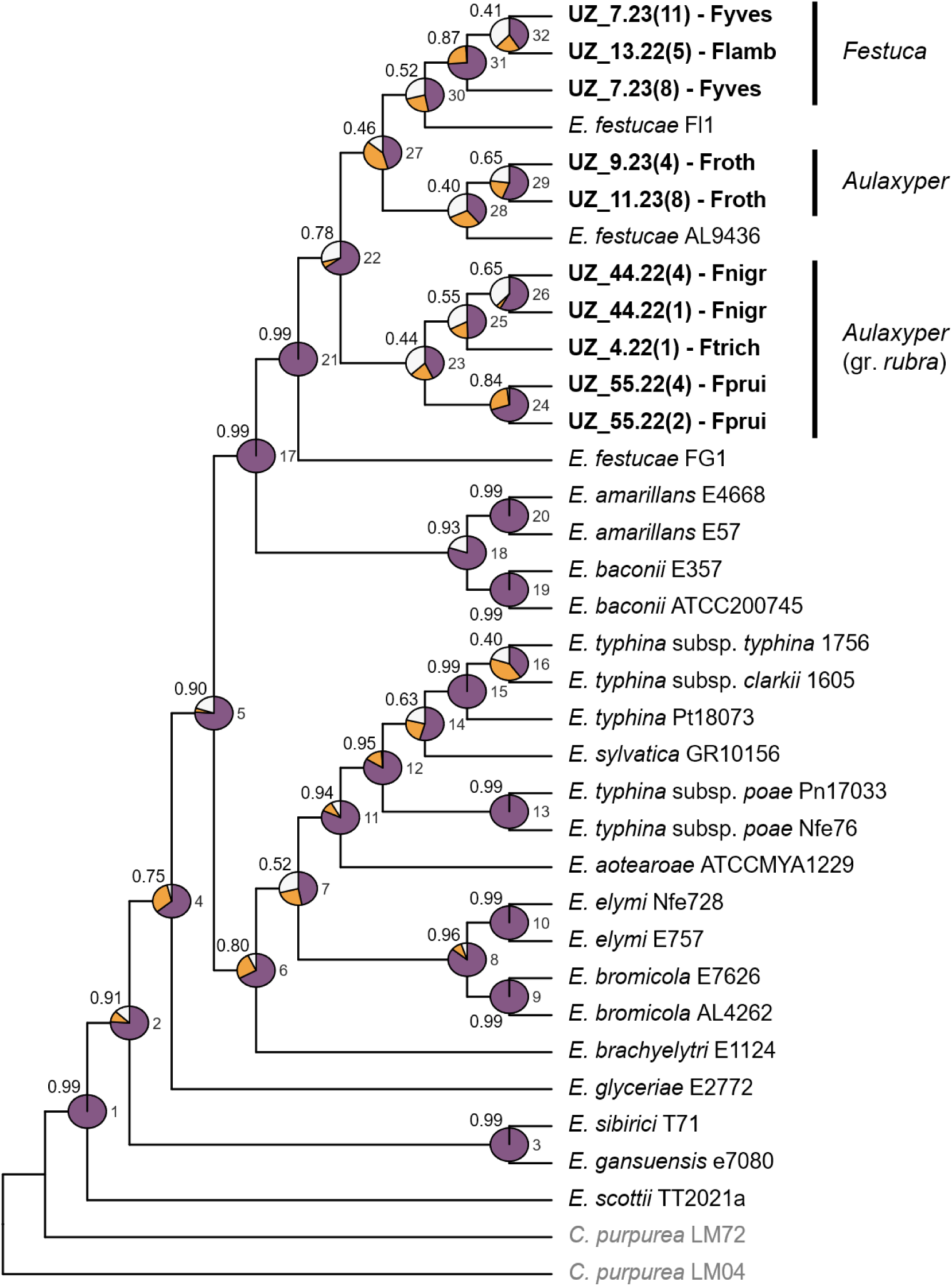
Multispecies coalescent (MSC) tree of *Epichloë festucae* constructed with ASTRAL III using as inputs the independent *actG*, *CalM*, ITS region, *tefA*, and *tubB* IQTREE2 ML trees (see also Fig. S2). Analyzed samples are highlighted in bold (Table S8), and external sequences of other *Epichloë* specimens and the outgroup *Claviceps purpurea* were obtained from NCBI (Sotomayor-Alge et al. 2025). The corresponding phylogenetic lineages of *Festuca* hosts are highlighted with vertical bars for the studied fungal samples, following Moreno-Aguilar et al. (2026). Values on branches correspond to posterior probability support (PPS). Quartet values indicate nodal support for the best tree (q1; purple) and the first (q2; orange) and second (q3; white) alternative topologies. Node numbers indicated in the phylogenetic tree and quartet values are listed in Table S7.

Single-locus ML phylogenies recurrently placed all newly sequenced isolates from fine-leaved *Festuca* holobionts within the *E. festucae* clade but differed markedly in their ability to resolve relationships within it (Figure S2). Among the five loci, *actG* provided the clearest holobiont-associated structure, recovering several isolate pairs or groups corresponding to their host species of origin, including Fprui, Fnigr, Froth and Fyves, with Flamb placed close to the Fyves lineage (Figure S2A). A weaker but partly comparable signal was observed for *CalM* and *tefA*, which recovered some host-associated pairs, particularly Fprui and Froth in *CalM* (Figure S2B) and Fnigr, Fprui and Fyves in *tefA* (Figure S2D). By contrast, ITS and *tubB* provided limited resolution within *E. festucae* and did not recover a consistent holobiont-associated topology (Figures S2C,E). Overall, no strongly supported conflict was detected among individual loci relative to the multilocus reconstructions.

### Alkaloid genetic and chemical profiles

PCR screening of alkaloid biosynthetic genes together with targeted alkaloid profiling of leaf extracts by UHPLC-QTOF/HRMS and GC–MS revealed substantial variation in alkaloid profiles among *Festuca*–*Epichloë* holobionts (Figure 4). Both inter- and intra-holobiont variability were observed across the four alkaloid families analyzed, although no more than two alkaloid families were chemically detected within the same holobiont. Overall, the alkaloid profiles of Farun (*E. coenophiala*) differed markedly from those associated with fine-leaved *E. festucae* holobionts. Within *E. festucae*, Froth and Fnigr5 showed particularly distinctive profiles, as both carried genes associated with pyrrolopyrazine, ergot alkaloid and indole-diterpene biosynthetic pathways, but chemically accumulated only pyrrolopyrazines and indole-diterpenes together with traces or absence of ergot alkaloids.

**Figure 4.**
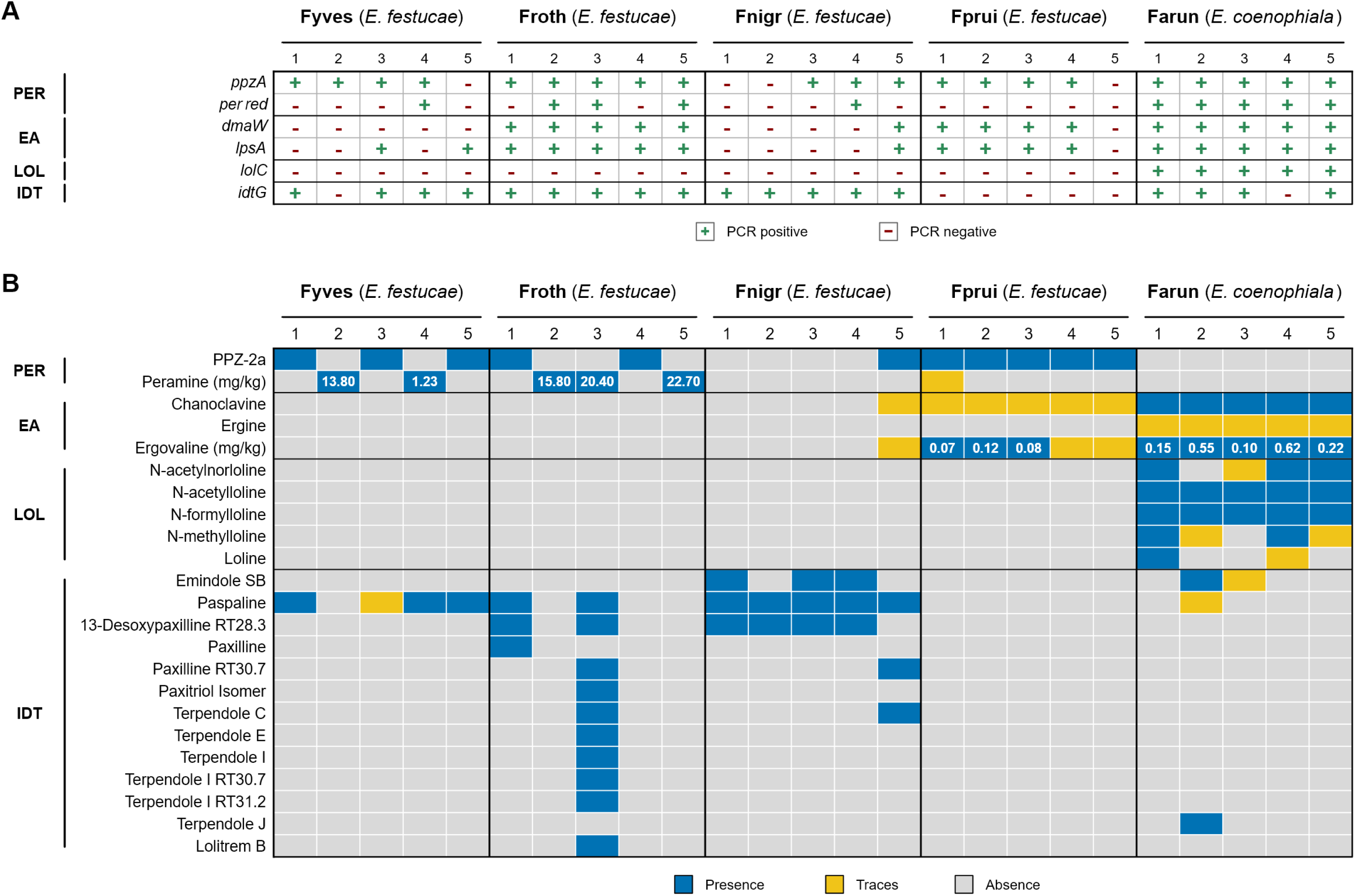
Alkaloid biosynthetic potential and chemical profiles across *Epichloë festucae* holobionts associated with different host species: *Festuca yvesii* (Fyves), *F. rothmaleri* (Froth), *F. nigrescens* (Fnigr) and *F. rubra* subsp. *pruinosa* (Fprui). The heteroploid endophyte *Epichloë coenophiala* (associated with *F. arundinacea* subsp. *arundinacea*; Farun) is also included. Within each holobiont, columns represent individuals (1–5). The four alkaloid families are considered: pyrrolopyrazines (PER), ergot alkaloids (EA), 1-aminopyrrolizidines (LOL) and indole-diterpenes (IDT). **(A)** PCR-based detection of key genes involved in alkaloid biosynthesis. **(B)** Qualitative and quantitative alkaloid profiles; the colors indicated on the chart represent the presence, traces, or absence of the respective alkaloids. Quantified concentrations (mg/kg) are shown for peramine and ergovaline when detected. Total 1-aminopyrrolizidines concentrations (mg/g) detected in Farun were: 3.52 (Farun1), 0.68 (Farun2), 0.66 (Farun3), 1.33 (Farun4), and 1.46 (Farun5). “RT” stands for isomers within a compound.

For pyrrolopyrazines (e.g., peramine), molecular detection targeted *ppzA* and its reductase domain (*per red*). All holobionts except Fprui harbored the *ppzA-1* allele together with the reductase domain associated with peramine biosynthesis (Figure 4A). This pattern was largely consistent with the chemical analyses, although Fyves2, Fnigr3, Fnigr4 and Fnigr5 lacked detectable peramine despite carrying the corresponding genes, whereas Farun uniformly carried the wild-type allele (*ppzA-1*) but showed no detectable peramine production. The only apparent inconsistency involved Froth2, likely reflecting a false-negative PCR result given the detection of peramine despite the apparent absence of the reductase domain (Figure 4B). When intermediate compounds accumulated, pyrrolopyrazine-1,4-dione cyclo prolyl-arginine (PPZ-1,4-dione 2a in Berry et al. 2019) was consistently detected, whereas other pyrrolopyrazine-1,4-diones (PPZ-1,4-dione 2b and PPZ- 1,4-dione 2c in Berry et al. 2019) were absent. Peramine concentrations ranged from 1.23 to 13.81 mg/kg in Fyves and from 15.84 to 22.75 mg/kg in Froth (Figure 4B), values matching or exceeding concentrations associated with insect deterrence (∼2 mg/kg; Vikuk et al., 2020; Realini et al., 2024).

Ergot alkaloid biosynthetic potential was assessed through PCR detection of *dmaW* and the non-ribosomal peptide synthetase gene *lpsA* involved in late pathway steps leading to ergovaline. These genes were repeatedly detected in Froth and Farun, present in most Fprui holobionts, and largely absent from Fyves and Fnigr (Figure 4A). Chemical analyses broadly mirrored these patterns, although no ergot alkaloids were chemically detected in Froth despite the presence of the biosynthetic genes. Ergovaline concentrations ranged from 0.10 to 0.62 mg/kg in Farun and from traces to 0.12 mg/kg in Fprui (Figure 4B), values within the range reported for endophyte-infected pastures associated with physiological effects in grazing mammals (Nicol and Klotz, 2016; Caradus et al., 2020; Klotz, 2022). In Farun, chanoclavine was additionally detected together with ergovaline. By contrast, ergovaline was not detected in Fyves and detected only as traces in Fnigr5. Interestingly, Fprui5 tested negative for ergot alkaloid genes despite exhibiting trace alkaloid accumulation.

For 1-aminopyrrolizidines, molecular detection targeted the early-stage *lolC* gene. This marker was detected exclusively in Farun and was consistently absent from all *E. festucae* holobionts (Figure 4A), fully consistent with the chemical analyses. Accordingly, 1- aminopyrrolizidines were detected only in Farun, including N-acetylnorloline, N- acetylloline, N-formylloline, N-methylloline and loline (Figure 4B), whereas the acetylated intermediate AcAP was not detected in any sample. Total 1-aminopyrrolizidine concentrations, calculated as the sum of these compounds, ranged from 0.66 to 3.52 mg/g and fell within the range commonly reported for alkaloid-producing *Epichloë* symbioses associated with strong insecticidal activity in grass–endophyte systems (Bush et al., 1997; Soto-Barajas et al., 2019; Cagnano et al., 2019; Realini et al., 2024).

Indole-diterpene biosynthetic potential was evaluated through PCR detection of the early-stage *idtG* gene. This marker was absent only in Fprui and present in all remaining holobionts (Figure 4A), a pattern broadly consistent with the qualitative chemical analyses for the indole-diterpene compounds listed in Figure 4B. Paxilline isomers RT 21.2 and RT 31.2 were screened but not detected in any holobiont. Specifically, paspaline was found in holobiont Fyves, paspaline, 13-desoxypaxilline isomer RT 28.3, paxiline, paxiline isomer RT 30.7, paxitriol isomer and terpendoles C, E, and I (including isomers RT 30.7 and RT31.2) in Froth, emindole SB, paspaline, 13-desoxypaxilline isomer RT 28.3, paxiline isomer RT 30.7, and terpendole C in Fnigr, and emindole SB, terpendole, and some traces of paspaline in Farun. In this case, individuals within holobionts displayed markedly different chemical profiles (e.g., Froth and Fnigr; Figure 4B). Although lolitrem B is the most extensively studied compound in this family, it was detected only in Froth3.

Overall, alkaloid profiling revealed marked differences between *Epichloë festucae* and *E*. *coenophiala* and substantial host-associated variation within *E. festucae* (Figure 4). Among the fine-leaved host species (*E. festucae*), ergot alkaloid production was largely restricted to Fprui, whereas Fnigr was characterized by the absence of detectable peramine and ergovaline despite consistent indole-diterpene production. Within this holobiont, the individual Fnigr5 displayed a distinct alkaloid profile relative to the remaining Fnigr samples. Fyves and Froth exhibited broadly comparable chemical profiles despite differing in their inferred biosynthetic gene complements.

The distinct alkaloid profiles identified here, many of them associated with particular host species, are of considerable interest for endophyte-mediated improvement of turf and forage grasses, where specific alkaloid phenotypes are desirable (Johnson et al., 2013). Because naturally occurring *E. festucae* chemotypes have never been comprehensively analyzed across Loliinae hosts, we compiled the genotypic and chemotypic profiles reported to date (Table 3). The survey revealed that only a limited number of genotype–chemotype combinations have been reported for naturally occurring *E. festucae* associations. Accordingly, the isolates characterized in the present study encompass a substantial proportion of the profiles described to date for fine-leaved Loliinae hosts. Furthermore, chemotypic characterization extended beyond the marker compounds routinely examined in previous reports (i.e., peramine, ergovaline and lolitrem B), incorporating a broader range of metabolites within each alkaloid class and revealing previously unreported genotype– chemotype combinations.

**Table 3.**
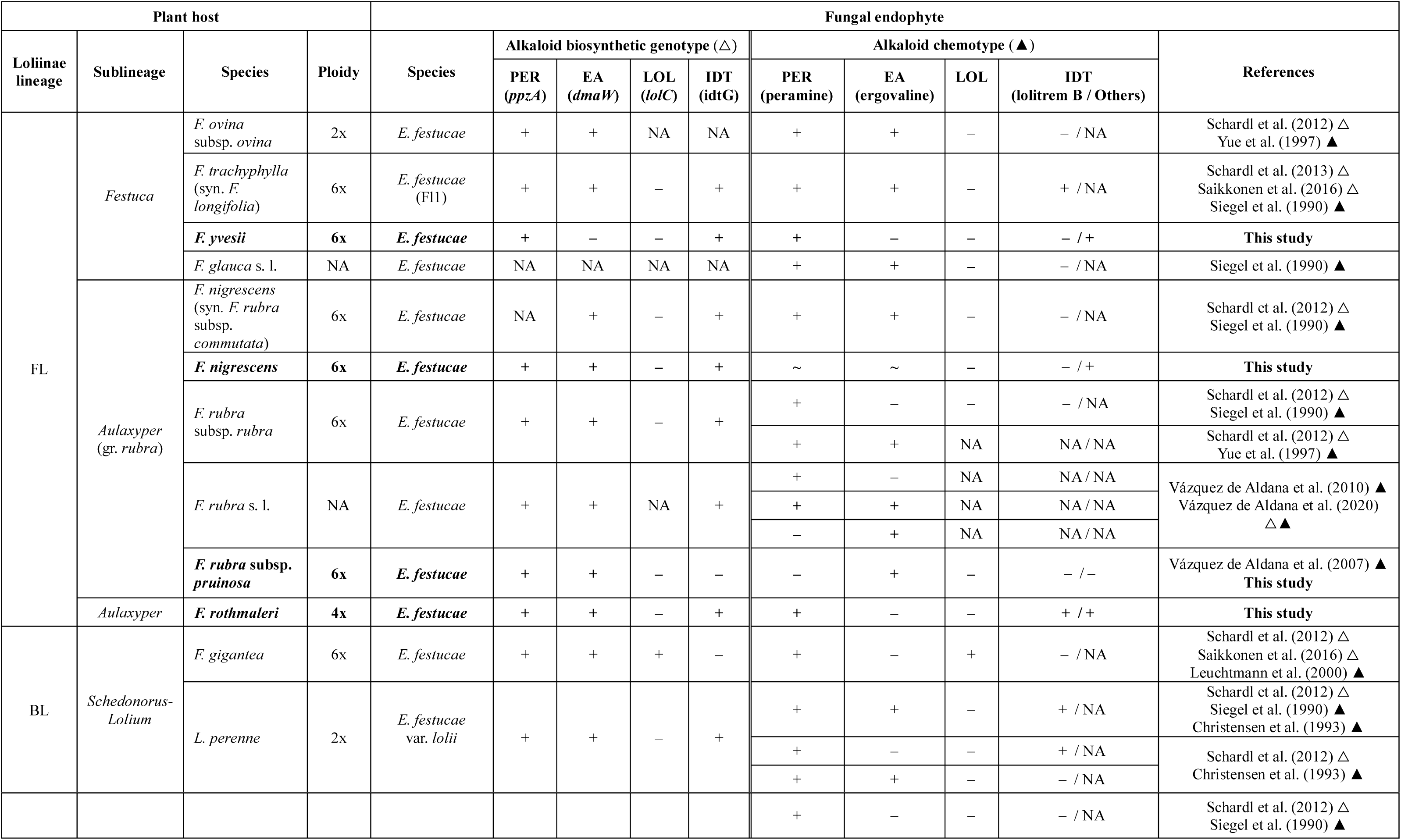
Genotypic and chemotypic alkaloid profiles of naturally occurring *Epichloë festucae* associated with fine-leaved (FL) and broad-leaved (BL) Loliinae hosts according to the phylogenomic framework of Moreno-Aguilar et al. (2026). Ploidy levels were inferred from Šmarda et al. (2008) and Garnatje et al. (2023) when not reported in the original reference(s) or in Moreno-Aguilar et al. (2026). The four alkaloid classes described in *Epichloë* are included: pyrrolopyrazines (PER), ergot alkaloids (EA), 1-aminopyrrolizidines (LOL) and indole-diterpenes (IDT). Genotypic profiles are based on the presence (+) or absence (–) of the alkaloid biosynthetic genes *ppzA* (PER), *dmaW* (EA), *lolC* (LOL) and *idtG* (IDT). Chemotypic profiles indicate the presence (+) or absence (–) of the corresponding alkaloid classes. In the consulted references, chemotypes were determined based on the detection of peramine (PER), ergovaline (EA) and lolitrem B (IDT). "NA" indicates that the corresponding variable was not analyzed, whereas "∼" indicates the detection of trace amounts of compounds within the corresponding alkaloid class. Profiles from this study are highlighted in bold.

## DISCUSSION

This study reveals two complementary signatures of host association across naturally occurring *Festuca–Epichloë* symbioses. First, multiple independent lines of biological evidence consistently differentiate *Epichloë festucae* strains according to host identity despite the broad geographic and ecological diversity of the sampled systems, indicating pervasive host-associated differentiation. Second, phylogenetic analyses recover a broader evolutionary pattern paralleling host phylogeny and reflecting a deeper historical signal. Together, these findings indicate that host identity consistently structures biological variation within *E. festucae*, whereas phylogenetic relationships preserve the deeper historical signature of host diversification.

### Ecological and evolutionary context of the studied *Festuca*–*Epichloë* symbioses

The *Festuca* species examined in this study encompass much of the ecological diversity of the genus in the Iberian Peninsula (Catalán, 2006; Devesa et al., 2020). Although *Festuca* exhibits extensive ploidy variation (Šmarda et al., 2008; Garnatje et al., 2023), the host species analyzed here are predominantly hexaploid while spanning two subgenera and three sections with distinct genome size patterns that reflect the evolutionary diversification within the genus (Table 2; Moreno-Aguilar et al., 2022b; Moreno-Aguilar et al., 2026). Thus, these host species provide an appropriate framework for assessing whether biological variation in *E. festucae* is consistently associated with host identity across evolutionarily divergent *Festuca* lineages and ecological settings.

*Epichloë* incidence varied widely among the studied host species and populations, ranging from complete absence (*F. ampla*, *F. elegans* and *F. paniculata*) or very low incidence (*F. trichophylla* and *F. lambinonii*) to frequencies exceeding 60% in *F. rubra* subsp. *pruinosa*. This variability agrees with previous reports of heterogeneous infection frequencies among host species (Siegel et al., 1984; Zabalgogeazcoa et al., 2003; Wäli et al., 2007; Iannone et al., 2009; Gundel et al., 2014) and among populations within the same species (Zabalgogeazcoa et al., 1999; Zabalgogeazcoa et al., 2006; Du et al., 2024; Sotomayor-Alge et al., 2025), and has generally been attributed to differences in host compatibility, transmission efficiency, fitness effects and local selective pressures (Clay and Schardl, 2002; Afkhami and Rudgers, 2008; Rudgers et al., 2009; Saikkonen et al., 2010; Dirihan et al., 2016; Vázquez de Aldana et al., 2026). Thus, variation in *Epichloë* incidence provides an ecological context for interpreting the pervasive host-associated differentiation revealed in the studied populations.

Consistent with its broad ecological distribution, *Epichloë festucae* has been reported from numerous Loliinae grasses occupying diverse environments (Schardl et al., 2013; Leuchtmann et al., 2014). However, relatively few naturally occurring host associations have been investigated using an integrative approach. The present study expands the known host range of *E. festucae* by characterizing novel associations with *Festuca trichophylla*, *F. lambinonii* and *F. yvesii*, while further assessing previously reported symbioses with *F. nigrescens* (Naffaa et al., 1998), *F. rubra* subsp. *pruinosa* (Zabalgogeazcoa et al., 2006; Pereira et al., 2019; Vázquez de Aldana et al., 2026), and *F. rothmaleri* (Sotomayor-Alge et al., 2025). Taken together, these host species encompass a broad ecological gradient (Table 1), providing a robust framework for assessing whether host-associated patterns remain consistent across contrasting environments and selective regimes (Rodríguez et al., 2009; Saikkonen et al., 2010; Sotomayor-Alge et al., 2025). Indeed, contrasting infection frequencies among host species occupying similar habitats—for example, the calcareous subalpine species of *F.* sect. *Festuca*, ranging from very low incidence in *F. lambinonii* to more than 60% in *F. yvesii*—suggest that host-related biological factors contribute to the establishment and persistence of these symbioses independently of local environmental conditions, supporting the interpretation that the patterns documented here are primarily associated with host identity.

### Host-associated phenotypic variation in *Epichloë festucae*

Variation in mating-type composition provides further evidence of differentiation among *E. festucae* populations associated with diverse host species. Although the present study was not designed as a population genetic survey, the recurrent occurrence of single or coexisting mating-type idiomorphs suggests contrasting reproductive scenarios among host-associated populations. Within the reproductive framework proposed for *Epichloë* (White, 1988; Tadych et al., 2014; Bultman et al., 2022), the mating-type compositions observed among the studied host populations are broadly consistent with distinct reproductive strategies. Populations in which only a single mating type was detected are compatible with predominantly clonal, vertically transmitted associations, whereas the coexistence of both idiomorphs in *F. yvesii* and *F. nigrescens* is compatible with mixed reproductive systems involving sexual reproduction and vertical transmission. However, the absence of stromata in *F. yvesii* despite the presence of both mating types, along with occasional stroma formation in *F. nigrescens*, demonstrates that mating-type composition alone is insufficient to determine expression of the sexual stage. Similar observations were reported for *F. rothmaleri* (Sotomayor-Alge et al., 2025) and are consistent with previous reports describing the infrequent occurrence of stromata in natural *E. festucae* populations (Leuchtmann et al., 1994; Zabalgogeazcoa et al., 2006), indicating that successful expression of the sexual stage requires additional biological and ecological factors beyond mating-type compatibility (White, 1988; Tadych et al., 2014). These observations suggest that differences in reproductive potential among these *E. festucae* strains may influence opportunities for recombination and gene flow, providing a plausible mechanism underlying the pervasive host-associated differentiation recovered across the independent biological datasets analyzed here, consistent with the evolutionary framework proposed by White (1988).

At the phenotypic level, the substantial morphological variability observed among *E. festucae* strains is consistent with the developmental plasticity of filamentous fungi, including *Epichloë* species (Steinberg et al., 2017). Despite this variability and the limited number of isolates analyzed, both linear mixed models and multivariate analyses revealed a host-associated structure of the asexual phenotype, evidencing that differences remain detectable despite substantial within-isolate variation (Figure 1; Table S1). Similar patterns have previously been reported in *Epichloë*, where both morphological and developmental traits differ among *Epichloë* strains associated with different host lineages (e.g., Du et al., 2024). Importantly, host-associated differentiation emerged from the combined contribution of morphological traits rather than from a single diagnostic character. Conidial area showed the strongest differentiation in the univariate analyses, whereas conidiogenous cell length contributed prominently to the multivariate separation of the studied *E. festucae* strains. Conversely, although conidial width exhibited comparatively weaker host-associated differentiation in the univariate analyses, it contributed strongly to multivariate separation, highlighting the complementary information captured by individual and multivariate approaches. Together, these results suggest that different components of the asexual phenotype may reflect distinct developmental constraints. In particular, the prominent contribution of conidiogenous cell length to multivariate differentiation is consistent with the polarized growth of filamentous fungi, in which elongation is generally more developmentally labile than radial expansion (Christensen et al., 2008; Harris, 2010; Steinberg et al., 2017). Nevertheless, the overall range of morphological values remained within that previously reported for *E. festucae* (Leuchtmann et al., 1994; Zabalgogeazcoa et al., 2006; Sotomayor-Alge et al., 2025), indicating that host-associated differentiation occurs within the normal intraspecific phenotypic limits of the species. By contrast, *E. coenophiala* was clearly differentiated from all *E. festucae* strains, consistent with its heteroploid origin and previous morphological descriptions (Leuchtmann et al., 2014).

This phenotypic differentiation was also reflected in the vegetative growth, with isolates from *F. nigrescens* exhibiting the fastest growth rates and those from *F. rubra* subsp. *pruinosa* the slowest, whereas *E. coenophiala* uniformly displayed markedly reduced growth relative to all *E. festucae* strains (Figure 2; Table S6). Overall, the congruence among reproductive traits, microscopic morphology and vegetative growth indicates that phenotypic differentiation in natural *E. festucae* populations is pervasive rather than trait-specific, reinforcing the view that host identity consistently structures phenotypic variation within the species.

### Genetic signatures of host-associated diversification in *Epichloë festucae*

At the phylogenetic level, genetic patterns were broadly consistent with the host-associated differentiation recovered across the phenotypic datasets. Multilocus analyses based on five nuclear loci (*actG*, *CalM*, ITS region, *tefA* and *tubB*) confidently assigned all isolates to the *Epichloë festucae* clade, while revealing two complementary levels of genetic structure. First, both phylogenetic approaches recurrently grouped isolates according to the phylogenetic lineages of their host species, with most isolate pairs from the same holobiont clustering together despite the limited number of loci analyzed (Figure 3; Figure S3). This recurrent clustering mirrors the host-associated differentiation inferred from morphology, vegetative growth and alkaloid profiles, suggesting that host identity also structures genetic variation within *E. festucae*.

Second, the multispecies coalescent reconstruction (MSC; Figure 3) resolved a broader evolutionary pattern that paralleled the phylogenetic relationships among the corresponding *Festuca* hosts (Inda et al., 2008; Minaya et al., 2017; Moreno-Aguilar et al., 2026). In particular, Flamb and Fyves formed a monophyletic lineage similar to that of their *Festuca* sect. *Festuca* hosts, while Froth separated from the clade comprising Fnigr, Ftrich and Fprui within the *E. festucae* “*F*. sect. *Aulaxyper*” subclade, reflecting the similar relationships reported for their *Festuca* sect. *Aulaxyper* hosts, which form a grade of *F. rothmaleri* and the *F. rubra*-aggregate clade (which includes *F. nigrescens*, *F. rubra* subsp. *pruinosa*, and *F. trichophylla*). By contrast, the concatenated maximum-likelihood analysis (ML; Figure S2) recovered stronger support for individual lineages associated with host identity but showed weaker correspondence with host phylogeny, indicating that the two approaches capture complementary aspects of evolutionary history. This difference likely reflects the ability of the MSC approach to accommodate incomplete lineage sorting (ILS), providing a more robust phylogenetic hypothesis when sequence divergence among loci is limited. Accordingly, the MSC reconstruction supports the divergence of *E. festucae* strains associated with the *F. sect. Festuca* clade (Flamb, Fyves), *F. rothmaleri* (Froth), and the typical *F. sect. Aulaxyper* clade (the *F. rubra* aggregate, including Fnigr, Ftrich and Fprui). A similar pattern has previously been proposed for host-specialized populations of the *E. typhina* complex, where strong genetic differentiation among host-associated lineages occurs despite incomplete phylogenetic concordance and occasional host shifts, supporting the hypothesis that prolonged host specialization may promote cryptic evolutionary divergence without requiring strict host–endophyte codiversification (Schardl et al., 2006). Consistent with this interpretation, the partial incongruence observed among loci and between phylogenetic approaches indicates that the evolutionary history of *E. festucae* is unlikely to reflect strict host–endophyte codiversification. Rather, our results fit a broader evolutionary scenario in which host jumps, host range expansions and occasional gene flow obscure, but do not erase, signals of long-term host–endophyte codiversification (Moon et al., 2002; Schardl et al., 2008; Tadych et al., 2014; Saikkonen et al., 2016; Thines, 2019, Catalán et al., 2025), leaving only a partial imprint of host evolutionary history on endophyte phylogeny. The predominantly vertical transmission, together with the infrequent expression of sexual reproduction in natural populations (Leuchtmann et al., 1994), may, however, facilitate the persistence of host-associated lineages observed in this study (Clay and Schardl, 2002; Saikkonen et al., 2004). Interestingly, several of the least-supported relationships in the MSC tree involved host populations in which both mating types were detected (Fyves, Froth and Fnigr), whereas Fprui, characterized by a single mating type, formed a cohesive lineage across both phylogenetic multilocus reconstructions. Although this observation remains tentative given the limited sampling available, it is compatible with the possibility that differences in opportunities for recombination contribute to variation in phylogenetic cohesion among the studied *E. festucae* strains.

Beyond phylogenetic relationships, genome size estimates provided an additional perspective on genetic variation. Haploid *E. festucae* strains exhibited remarkably limited variation in nuclear DNA content despite their contrasting phenotypic and functional characteristics and their association with hosts differing in ploidy level (Table 2). These results agree with previous flow-cytometric estimates (Sotomayor-Alge et al., 2025) and suggest that host-associated differentiation is unlikely to arise from large-scale variation in genome size but instead reflects finer-scale genomic or regulatory differences that warrant comparative genomic and transcriptomic analyses. By contrast, *E. coenophiala* displayed an approximately three-fold larger genome, consistent with its well-established triheteroploid origin (Moon et al., 2004).

Taken together, these findings reveal two complementary genetic signatures in natural *E. festucae* populations despite remarkably stable genome size across host lineages. At shallower evolutionary scales, isolates clustered according to host identity, whereas at deeper evolutionary scales the MSC reconstruction retained a historical signal broadly reflecting host phylogeny. Higher-resolution phylogenomic datasets will be required to determine whether these *E. festucae* strains represent ongoing evolutionary divergence within the species (e.g., Oberhofer and Leuchtmann, 2012; Catalán et al., 2025).

### Functional diversification of alkaloid profiles in *Epichloë festucae*

Alkaloid profiles displayed the greatest functional diversity among all datasets analyzed in this study, highlighting alkaloid production as the most variable biological dimension examined (Figure 4). Unlike the phylogenetic relationships recovered from multilocus analyses, alkaloid profiles did not mirror the evolutionary structure of the three *Epihcloë festucae* lineages identified here (Figure 3). Instead, chemically similar profiles were identified in phylogenetically distant host-associated populations (e.g. Fyves and Froth), whereas closely related lineages within the *F. rubra* aggregate displayed contrasting alkaloid complements (e.g. Fnigr and Fprui). These observations indicate that alkaloid production represents a more dynamic functional dimension of this symbiosis than either morphology or phylogenetic relationships. Nevertheless, characteristic chemotypes repeatedly occurred within particular host-associated populations, indicating that host identity still contributes to shaping functional variation despite the influence of regulatory and ecological factors.

A second prominent pattern was the incomplete correspondence between alkaloid biosynthetic genotype and chemical expression. Although the genetic detection of biosynthetic pathways generally agreed with the presence of the corresponding alkaloid classes, several recurrent mismatches were observed, reinforcing previous evidence that the presence of biosynthetic genes alone does not necessarily predict alkaloid accumulation *in planta* (Schardl et al., 2013; Panaccione et al., 2014; Bastias et al., 2017). Across all host-associated populations, no *E. festucae* holobiont consistently accumulated more than two alkaloid classes despite frequently carrying markers for additional biosynthetic pathways (Figure 4), reinforcing that alkaloid production is constrained by regulatory processes, pathway interactions or ecological trade-offs rather than by biosynthetic potential alone (Panaccione et al., 2014 Berry et al., 2015; Young et al., 2015; Vázquez de Aldana et al., 2020).

The Farun holobiont illustrates this complexity particularly well: despite apparently possessing the genetic machinery for peramine biosynthesis, it did not accumulate detectable peramine while simultaneously producing high concentrations of 1-aminopyrrolizidines together with ergot alkaloids. Although the mechanisms remain unresolved, these observations suggest that different *Epichloë* lineages may rely on alternative combinations of defensive metabolites, potentially reducing dependence on particular alkaloid classes. These general patterns became evident when individual alkaloid families were examined (Figure 4). Peramine production was largely associated with the presence of the reductase domain (Ekanayake et al., 2017; Berry et al., 2019; Hettiarachchige et al., 2019), although several Fnigr individuals and the Farun holobiont represented notable exceptions. A similar genotype–chemotype mismatch was observed for ergovaline. Whereas Farun and Fprui accumulated the alkaloid, Froth uniformly carried the corresponding biosynthetic markers without detectable ergot alkaloid production, indicating that completion of the pathway depends on additional genetic components or regulatory mechanisms beyond the markers analyzed in this study (Saikkonen et al., 2010; Schardl et al., 2013, Vázquez de Aldana et al., 2020). The considerable genetic diversity previously documented within natural *E. festucae* populations may further contribute to this variation in alkaloid expression (von Cräutlein et al., 2021). By contrast, 1-aminopyrrolizidines (AcAP and loline alkaloids) exhibited complete agreement between genotype and chemotype, being detected exclusively in the Farun holobiont. This pattern agrees with previous studies showing that the LOL biosynthetic cluster is restricted to particular *Epichloë* lineages and reflects its evolutionary history (Schardl et al., 2007; Schardl et al., 2013; Pan et al., 2014). As summarized in Table 3, these alkaloids have been reported from broad-leaved (*Schedonorus*–*Lolium*) associations, particularly those involving *Festuca gigantea* and *Epichloë festucae*, whereas they have not been described in naturally occurring fine-leaved *Festuca* spp*.– Epichloë festucae* symbioses.

Indole-diterpenes showed a third intermediate scenario. The core biosynthetic pathway was genetically and chemically detected in several host-associated populations, but the complexity of the accumulated metabolites differed markedly among them (Figure 4). Fyves accumulated only paspaline, whereas Froth and Fnigr produced a broader suite of intermediates, including terpendoles, paxilline derivatives and emindole SB, suggesting differences not only in pathway presence but also in the extent to which biosynthesis proceeds towards downstream metabolites. This agrees with previous studies showing that indole-diterpene biosynthesis involves numerous genes and is strongly influenced by regulatory processes and host- and environment-dependent factors (May et al., 2008; Schardl et al., 2013; Guerre, 2016; Fuchs et al., 2017). Accordingly, detection of pathway intermediates supports the presence of a functional core pathway, although not necessarily the complete synthesis of the most complex end products (Young et al., 2009). In this study lolitrem B, the main indole-diterpene compound studied to date, was only detected in one sample (Froth3).

The ecological significance of these differences remains uncertain but is unlikely to be explained solely by phylogenetic relationships. Alkaloid production is known to respond to multiple interacting factors, including fungal genotype, host genotype, environmental conditions and both abiotic and biotic stress (Saikkonen et al., 2010; Schardl et al., 2013, Vázquez de Aldana et al., 2020). Consequently, the contrasting chemotypes revealed here probably reflect the combined effects of evolutionary divergence, host-associated regulation and local ecological conditions. This greater functional lability is consistent with the ecological role of alkaloids as defensive metabolites, whose expression may respond more rapidly to local selective pressures than neutral phylogenetic markers, thereby explaining their weaker correspondence with the deeper phylogenetic structure recovered by the MSC analysis.

Beyond their biological implications, the alkaloid profiles reported here substantially expand the currently documented chemotypic diversity of naturally occurring *Epichloë festucae* holobionts. Naturally occurring genotype–chemotype combinations remain poorly documented, particularly in fine-leaved Loliinae clades (*Festuca* sects. *Festuca* and *Aulaxyper*), whereas broad-leaved *Schedonorus*–*Lolium* associations have historically received considerably greater attention because of their importance in forage and turf breeding programmes (Johnson et al., 2013; Card et al., 2021; Card et al., 2024). The synthesis presented in Table 3 highlights how fragmentary current knowledge remains for naturally occurring fine-leaved *Festuca–Epichloë festucae* associations. Integrative studies combining biosynthetic genotyping with metabolite profiling are still uncommon in these natural systems, making the *E. festucae* strains characterized here a substantial proportion of the currently documented *E. festucae* chemotypes from fine-leaved *Festuca* hosts. While the pyrrolopyrazine/ergot alkaloid chemotype is the predominant profile among fine-leaved Loliinae species, our findings expand this pattern by identifying several species with a pyrrolopyrazine/indole diterpene chemotype, a relatively rare combination. By combining biosynthetic genotyping with metabolite profiling beyond the traditional marker compounds (peramine, ergovaline and lolitrem B), this study identifies previously unreported genotype– chemotype combinations and broadens the known functional diversity of natural *Festuca*– *Epichloë* symbioses. Our results further support that this variation occurs not only among host-associated populations but also among individuals within populations, underscoring the remarkable plasticity of alkaloid expression. Together, these findings provide one of the most comprehensive characterizations of alkaloid diversity currently available for naturally occurring fine-leaved *Festuca–Epichloë* symbioses and establish a valuable foundation for future ecological, evolutionary and applied studies of grass–endophyte interactions.

### Towards a multidimensional understanding of host -associated diversification in *Epichloë festucae*

This study shows that host-associated differentiation in *Epichloë festucae* is a multidimensional phenomenon encompassing reproductive biology, phenotype, genetic structure and secondary metabolism, while expanding the known diversity of naturally occurring *Festuca–Epichloë* associations in the Iberian Peninsula. Collectively, these complementary datasets reveal two interconnected signatures of diversification: pervasive host-associated differentiation across biological dimensions and a deeper historical signal retained in phylogenetic relationships. Our findings demonstrate the value of integrating complementary biological approaches to understand diversification in naturally occurring plant–fungal symbioses and provide a foundation for future phylogenomic, ecological and functional studies of the *Festuca–Epichloë* system.

## MATERIALS AND METHODS

### Field sampling and host identification

Plant material was collected from natural ecosystems across the northern Iberian Peninsula (Spain). Specifically, *Festuca lambinonii*, *F. yvesii*, and *F. nigrescens* were sampled in the pre-Pyrenean and Pyrenean regions of Huesca province; *F. trichophylla* in the central Ebro river valley (Zaragoza province); *F. rothmaleri*, *F. ampla*, *F. elegans* and *F. paniculata* from the mountain ranges of southern Salamanca province (Sotomayor-Alge et al., 2025); and *F. rubra* subsp. *pruinosa* along the coastal ecosystems of the Atlantic coast of Galicia (Table 1). To minimize the likelihood of clonal sampling, individuals were collected at least 3 m apart from one another. Plants were transplanted into pots containing universal substrate (Blumenerde, Gramoflor) and maintained under comparable temperature (20–27 °C) and watering conditions (approximately 3 times per week depending on seasonal conditions) throughout the study. Between 10 and 30 individuals per species were processed, and voucher specimens were deposited at the herbarium of the High Polytechnic School of Huesca, University of Zaragoza (Spain). Taxonomic identification of *Festuca* taxa was conducted according to the main taxonomic keys for the genus (Al-Bermani et al., 1992; Devesa et al., 2020), using morphoanatomical traits for the characterization at species level (e.g., measurements of vegetative and reproductive structures and leaf cross sections). As an outgroup for both host and endophyte analyses, naturally occurring individuals of the broad-leaved species *F. arundinacea* subsp. *arundinacea* harboring *Epichloë coenophiala* were collected in the pre-Pyrenean region of Huesca and processed alongside the fine-leaved holobionts (Table 1).

### Endophyte detection, isolation and incidence

*Epichloë* endophytes were detected in above-ground plant tissues using both aniline blue staining and direct isolation (Florea et al., 2015). For fungal isolation, surface-sterilized fragments of floral stems or basal tillers were cultured on potato dextrose agar (PDA; Condalab) supplemented with chloramphenicol (25 µg/mL; PanReac AppliChem) to inhibit bacterial growth. Plant fragments were surface sterilized in 20% bleach for 10 min prior to cultivation. Emergent *Epichloë*-like colonies were subcultured on PDA and maintained at 22–25 °C in darkness for subsequent analyses. Endophyte incidence was estimated for each host population as the proportion of plants harboring the *Epichloë* endophyte relative to the total number of individuals analyzed.

### Endophyte mating-type composition

During field sampling, stromata were occasionally observed in *F. nigrescens*, whereas none were detected in the remaining host species, including *F. arundinacea*, whose hybrid endophyte is considered strictly asexual (Schardl et al., 2013; Leuchtmann et al., 2014). To assess the potential for sexual reproduction, PCR screenings of the *MTA* (*mtAC*; 785 bp) and *MTB* (*mtBA*; 215 bp) mating-type idiomorphs were performed following Florea et al., (2015) using total DNA extracted from leaf tissue. In populations where only one idiomorph was detected, all individuals available were processed. Primer sequences and PCR conditions are listed in Table S9.

### Cytogenetic profiling of *Festuca*–*Epichloë* holobionts

Genome size of *Festuca* hosts harboring *Epichloë* endophytes was estimated by flow cytometry (Ploidy Analyzer, Sysmex) using fresh mature leaf tissue, following the protocol of Doležel et al. (2007) with Otto I and Otto II buffers. Up to five individuals per species were analyzed, each with two technical replicates (n = 10). Measurements were retained only when the number of nuclei exceeded 5,000 and coefficients of variation (CVs) were below 3.5%. *Pisum sativum* ‘Ctirad’ (9.09 pg/2C) and *Secale cereale* ‘Daňkovské’ (16.19 pg/2C) were used as an internal standards (Doležel et al., 2007). Genome size categories were subsequently compared with published chromosome counts and polyploidy data in *Festuca* to infer ploidy levels (e.g., Šmarda et al., 2008; Garnatje et al., 2023).

Similarly, genome size of *Epichloë* isolates was estimated using the same flow cytometry platform following Sotomayor-Alge et al. (2025). In this case, up to three isolates per host species, each represented by two technical replicates (n = 6), were analyzed using *Colletotrichum acutatum* strain PT812 (68 Mb; ∼0.069 pg/1C) as internal standard (Talhinhas et al., 2017). Measurements were retained only when at least 5,000 nuclei were recorded and CVs remained below 10% (Bourne et al., 2014).

### Morphological and growth characterization of *Epichloë* strains

To assess morphological variation among endophytes with different host species, three fungal isolates per host species with three biological replicates each were considered whenever available. Microscopic preparations consisted of 1 mm² mycelial sections grown on water agar (WA; Bacteriological Agar, Condalab) for 3 weeks in darkness and subsequently mounted under coverslips. For each *Epichloë* isolate, 10 conidia (length, width, and area) and 10 conidiogenous cells (total length and basal width) were measured per replicate, resulting in up to 30 measurements per isolate. These assessments were conducted following the procedures described in Sotomayor-Alge et al., (2025), with minor modifications to the “*Epichloë conidia*” software to accommodate measurements of heteroploid *E. coenophiala* conidia.

Morphological differences among *Epichloë* specimens were then analyzed using linear-mixed effects models implemented in the R package lme4 (v2.0.1; Bates et al., 2015). For each trait, host species was included as a fixed effect, whereas *Epichloë* isolates and technical replicates nested within isolates were treated as random intercepts. Models were fitted using restricted maximum likelihood (REML) method, and the significance of host species effects was assessed using type III ANOVA with Kenward–Roger approximation (Kenward and Roger, 1997; Kuznetsova et al., 2017). Marginal and conditional R² values were calculated to estimate the variance explained by fixed effects alone and by the full model, respectively. Model assumptions were evaluated and, when minor deviations from normality or homoscedasticity were detected, additional sensitivity analyses were performed using log-transformed response variables.

Multivariate differences in morphological traits among host-associated strains were assessed using Euclidean distance-based PERMANOVA with 10,000 permutations, implemented in the R package *vegan* (v2.7.5; Oksanen et al., 2025). Distance matrices were generated from standardized morphological variables, and permutations were constrained within *Epichloë* isolates to account for non-independence among technical replicates. Pairwise PERMANOVA comparisons were adjusted for multiple testing using false discovery rate (FDR) correction (Benjamini and Hochberg, 1995), whereas homogeneity of multivariate dispersion was evaluated using the *betadisper* procedure. Patterns of multivariate morphological variation among *Epichloë* strains were further explored using principal component analysis (PCA) based on standardized independent variables, implemented in the R package *FactoMineR* (v2.14; Lê et al., 2008). Differentiation along the main ordination axes was evaluated using one-way ANOVAs on PC1 and PC2 scores followed by Tukey-adjusted pairwise comparisons. To ensure balanced statistical comparisons, main analyses were restricted to endophytes from host species represented by three isolates (i.e., *F. yvesii*, *F. rothmaleri, F. nigrescens*, and *F. rubra* subsp. *pruinosa*).

Growth rate (GR) was assessed for all *Epichloë* strains following Sotomayor-Alge et al. (2025), with minor modifications. Briefly, 2 mm² sections from two-week-old isolates were grown on PDA plates for 21 days in darkness at room temperature (22–25 °C), and colony diameters were measured weekly to estimate growth rates (mm/day). Whenever possible, three *Epichloë* isolates and three technical replicates per isolate were analyzed for each host species. Isolates from *F. rothmaleri* were re-cultivated alongside the newly obtained isolates to ensure comparability among measurements. Growth rate differences among host-associated strains were analyzed using linear mixed-effects models, with host species included as a fixed effect and *Epichloë* isolates nested within host species treated as random intercepts. Significance was assessed using Type III ANOVA, and post hoc pairwise comparisons were performed using Tukey-adjusted estimated marginal means with Kenward–Roger approximation.

### Molecular and phylogenetic analyses of *Epichloë* isolates

Five nuclear barcode loci, including the coding genes γ-actin (*actG*), calmodulin (*CalM*), translation elongation factor 1-α (*tefA*), and β-tubulin (*tubB*), together with the rDNA ITS region, were used for molecular characterizations. *Epichloë* isolates from two specimens per host species were grown on PDA plates covered with sterile cellophane disks for 7–10 days. Genomic DNA was extracted following the CTAB-based protocol described in Sotomayor- Alge et al. (2025), and DNA quality and concentration were assessed using a Biodrop spectrophotometer (µLite) and a Qubit 3.0 fluorometer. PCR amplifications were performed using KAPA Taq polymerase (Kapa Biosystems) following manufacturer recommendations. PCR products were purified with IllustraTM ExoProStarTM 1-Step (GE Healthcare, Life Sciences) and subjected to bidirectional Sanger sequencing at STAB VIDA (Caparica, Portugal). Primers and cycling programs are listed in Table S9. Because PCR cloning was not performed, multiple gene copies derived from the heteroploid genome of *E. coenophiala* could not be resolved, and this species was therefore excluded from phylogenetic analyses. Newly generated sequences were deposited in GenBank (Table S8).

Sequences were trimmed, aligned using MAFFT algorithm v7.490 (Katoh and Standley, 2013), and manually refined in Geneious Prime version 2026.0.2 (Biomatters Ltd, New Zealand). Reference and outgroup sequences were retrieved from NCBI as indicated in Sotomayor-Alge et al. (2025). Single-locus maximum likelihood phylogenetic trees were inferred using IQ-TREE 2 (Minh et al., 2020) with UltraFast Bootstrap (BS) support (Hoang et al., 2017). Branch lengths were transformed using x^¼^ for visualization purposes. Concatenated maximum likelihood and coalescent-based species trees were subsequently reconstructed using IQ-TREE 2 and ASTRAL-III (Zhang et al., 2018), respectively. The likelihood-based site concordance factors (sCFLs) were calculated for the concatenated ML tree (Mo et al., 2023), whereas posterior probabilities and quartet scores were estimated for the coalescent-based species tree following Sayyari and Mirarab (2016). All trees were visualized and formatted using adapted bash and R Markdown scripts from in Sotomayor- Alge et al. (2025).

### Alkaloid genotyping and chemical profiling

Alkaloid profiling was conducted in holobionts with sufficient *Epichloë* incidence to include five plants harboring the endophyte (E^+^) and one endophyte-free specimen (E^-^) plant per host association. Plants were maintained under comparable greenhouse conditions for eight months prior to sampling. Material was collected within the same day at similar phenological stages from multiple vegetative aerial tissues, immediately placed on ice to halt metabolic activity, and stored at −80 °C until lyophilization. Freeze-dried material was ground using a homogenizer (Biospec Scientifica mill) and subsequently used for both genotypic screening of alkaloid biosynthetic pathways and chemical characterization of alkaloid profiles.

For genotypic profiling, total DNA was extracted from 20 mg of ground plant material using the DNeasy Plant Mini Kit (Qiagen, Valencia, CA, USA) following the manufacturer recommendations. PCR screening targeted key genes involved in the four major *Epichloë* alkaloid biosynthetic pathways, using the primers and conditions described in Table S9. The primer pair *perT2* enables detection of the *ppzA* gene involved in pyrrolopyrazine biosynthesis, whereas *per red* targets the C-terminal reductase domain of *ppzA*, reporting gene integrity and thus the potential to produce peramine (Berry et al., 2015, Berry et al., 2019). Primers *dmaW* and *lpsA* are associated with the ergot alkaloid pathway, representing early and late steps in ergovaline biosynthesis, respectively. *lolC* serves as a marker for early steps in 1-aminopyrrolizidine biosynthesis, whereas *idtG* indicates the initial stages of indole-diterpene synthesis, including lolitrems (Schardl et al., 2013). *E. coenophiala* strains e19 and 4163 were used as positive controls, and PCR products were separated on 2% agarose gels in TBE buffer and visualized under UV transillumination (Bio-Rad).

Chemical analyses included qualitative and quantitative characterization of pyrrolopyrazines (e.g., peramine), ergot alkaloids (e.g., ergovaline), and 1-aminopyrrolizidines (i.e., loline alkaloids), whereas indole-diterpenes (e.g., lolitrems) were only assessed qualitatively. Pyrrolopyrazines (peramine and pyrrolopyrazine-1,4-diones) and ergot alkaloids (ergovaline, ergine and chanoclavine) qualitative detection was conducted following Nagabhyru et al. (2026) using UHPLC-QTOF mass spectrometry. Briefly, freeze dried finely ground plant tissue (25 mg) was extracted twice using an 80:20 methanol:water solution with 0.1% formic acid. Each extraction involved vortexing 10 min, sonication for 5 min and centrifugation at 16000 RCF for 10 min, after which supernatants were collected. The combined extracts were re-centrifuged to obtain a clear supernatant and transferred to clear glass vials. The chromatographic separation was performed using an Agilent 1290 Infinity II UHPLC with a Luna Omega C18 column, coupled to an Agilent 6546 QTOF mass spectrometer operating in positive mode. In this case, ergovaline and peramine hemisulfate (BenChem) and chanoclavine (a gift from Dr. Daniel G. Panaccione, West Virginia University) were used as external standards. For 1-aminopyrrolizidines, compounds were analyzed by GC–MS following Pan et al. (2014). Briefly, extraction was performed using 100 mg of ground lyophilized plant tissue in 1 mL of chloroform under alkaline conditions, achieved by the addition of 100 µL of 1N NaOH. Samples were analyzed by gas chromatography–mass spectrometry (GC–MS) using an Agilent 7890 gas chromatograph coupled to an Agilent 5975 mass spectrometer (Agilent Technologies, Santa Clara, CA, USA). Here, analytical grade quinoline (Sigma Aldrich, St. Louis, MO) was used as an internal standard and pure N-formylloline (NFL, purified from tall fescue seeds following procedure described in (Bacetty et al., 2009) was used as the reference standard for identification and quantification of major 1-aminopyrrolizidine alkaloids. Indole-diterpenes were qualitatively assessed by HPLC-HRMS following Rasmussen et al. (2012) and Lee et al. (2017) including up to 15 different compounds. Briefly, 50 mg of ground plant material was extracted with isopropanol, followed by centrifugation and filtration. Extracts were analyzed by HPLC-HRMS using a Betasil C18 reversed-phase column. Separation was achieved with a 0.1% formic acid– acetonitrile gradient (80:20 to 100% acetonitrile over 40 min) at 0.3 mL/min on an Ultimate 3000 HPLC coupled to an Exactive Plus Orbitrap HRMS (HESI). Compounds were identified by reconstructed ion chromatograms using calculated [M+H]⁺ exact masses with a mass tolerance of 10 ppm. Previously characterized endophyte infected *Ipomoea asarifolia* and *Ipomoea muelleri* seed materials were used as qualitative external standards (Lee et al., 2017).

Quantitative analyses of ergovaline and peramine were performed following Vassiliadis et al. (2023) with minor modifications. Briefly, 20 mg of ground lyophilized plant material was extracted twice with 1 mL of 80% methanol. After centrifugation, the two supernatants were combined, and the pooled extract was evaporated to dryness using a Savant SpeedVac concentrator. The resulting residue was resuspended in 80% methanol and filtered through a 0.22 µm nylon membrane prior to chromatographic analysis. Samples were analyzed on a UHPLC system (Agilent 1290 Infinity II) coupled to a quadrupole time of flight mass spectrometer (QTOF, Agilent G6546A), using a Zorbax Eclipse Plus C18 HD column maintained at 30 °C. The mobile phase consisted of 0.1% aqueous formic acid and acetonitrile. Ergovaline and peramine hemisulfate (BenChem) were used as external standards for quantification. For 1-aminopyrrolizidines, total alkaloid concentration was quantified relative to the NFL standard according to Pan et al. (2014).

## Data statement

The sequence data generated for this project is available in the NCBI GenBank database (http://www.ncbi.nlm.nih.gov) under the accession numbers summarized in Table S8. Sampling spots have been uploaded to GBIF database. The data and scripts underpinning the analyses were deposited in our GitHub repository (https://github.com/Bioflora/FestucaEpichloeComparativeStudy).

## Acknowledgements

We thank Pedro Talhinhas for kindly providing the fungal primary standard *Colletotrichum acutatum* strain PT812, Manuel Pimentel for sampling assistance, and Rebeca Alonso Nieto, Virginia González Blanco and Beatriz Larruy García for technical assistance.

## Author contributions

PC and ASA designed the study. ASA, LAI, BRV, IZ and PC collected plant specimens. ASA, PN, BRV, STL, LAI, CLS and PC conducted and supervised the analyses. ASA wrote the original draft, and all authors reviewed and wrote the final manuscript.

## Supplementary figures

**Figure S1.**
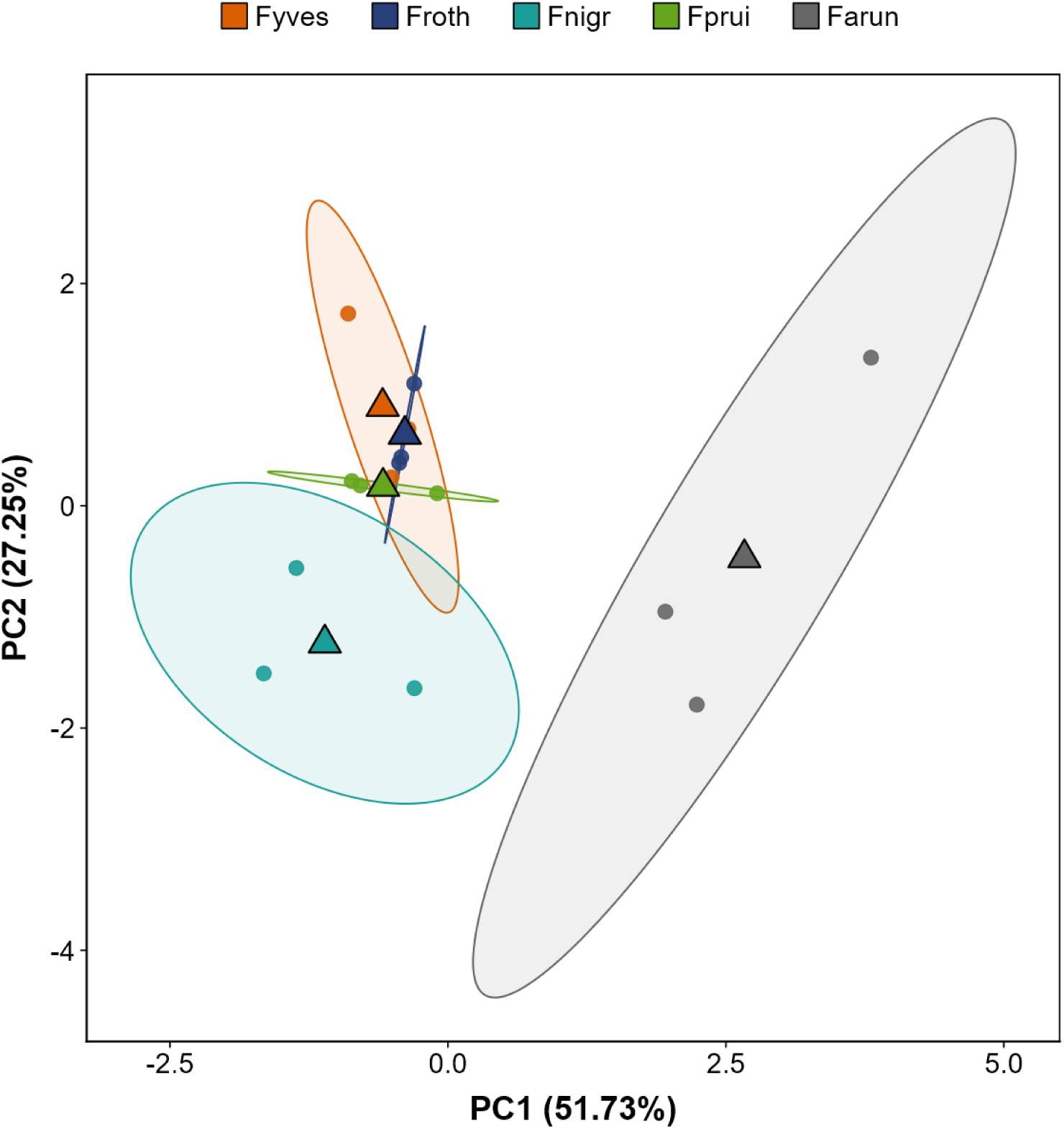
Principal component analysis based on independent asexual reproductive structures (conidia and conidiogenous cells) of *Epichloë* isolates associated with different *Festuca* host species. Holobionts abbreviations designated as follows: *Festuca yvesii* (Fyves), *F. rothmaleri* (Froth), *F. nigrescens* (Fnigr), *F. rubra* subsp. *pruinosa* (Fprui), and *F. arundinacea* subsp. *arundinacea* (Farun). Only species with three isolates were considered. Points represent individual isolates, colored according to holobiont identity, as shown in the chart. Ellipses indicate the dispersion of each group (95% CI), and triangles represent group centroids. See also Table S5.

**Figure S2.**
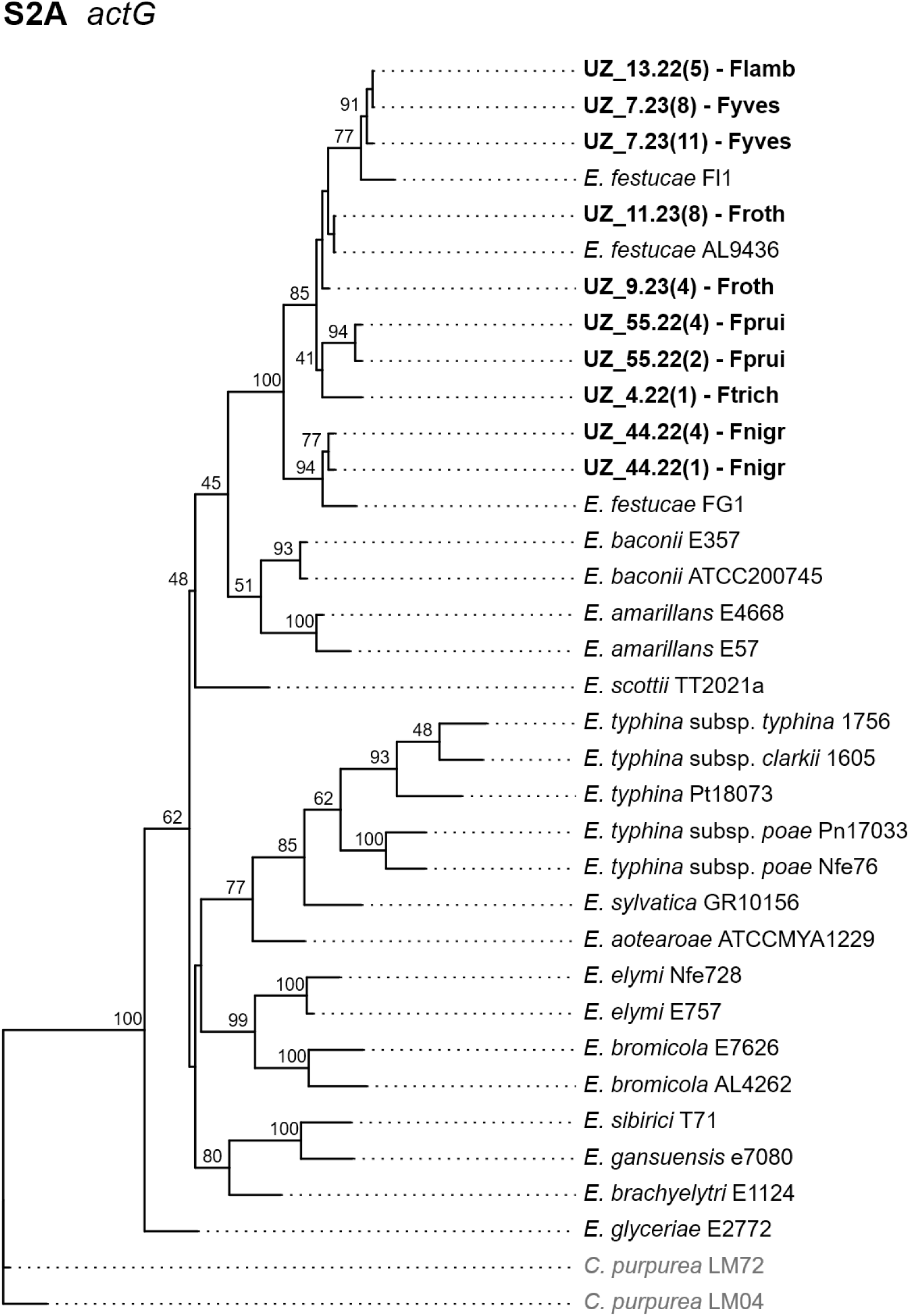
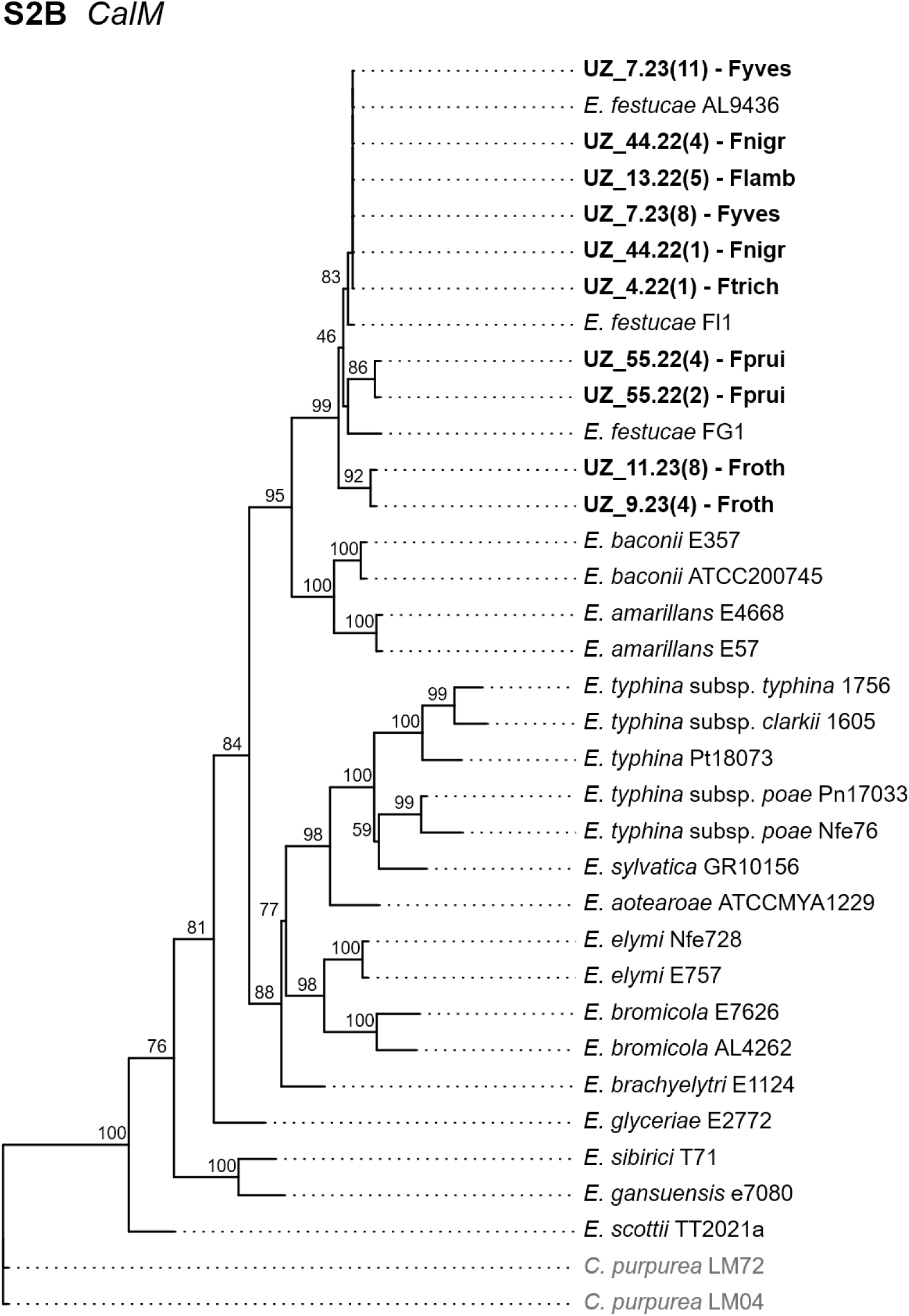
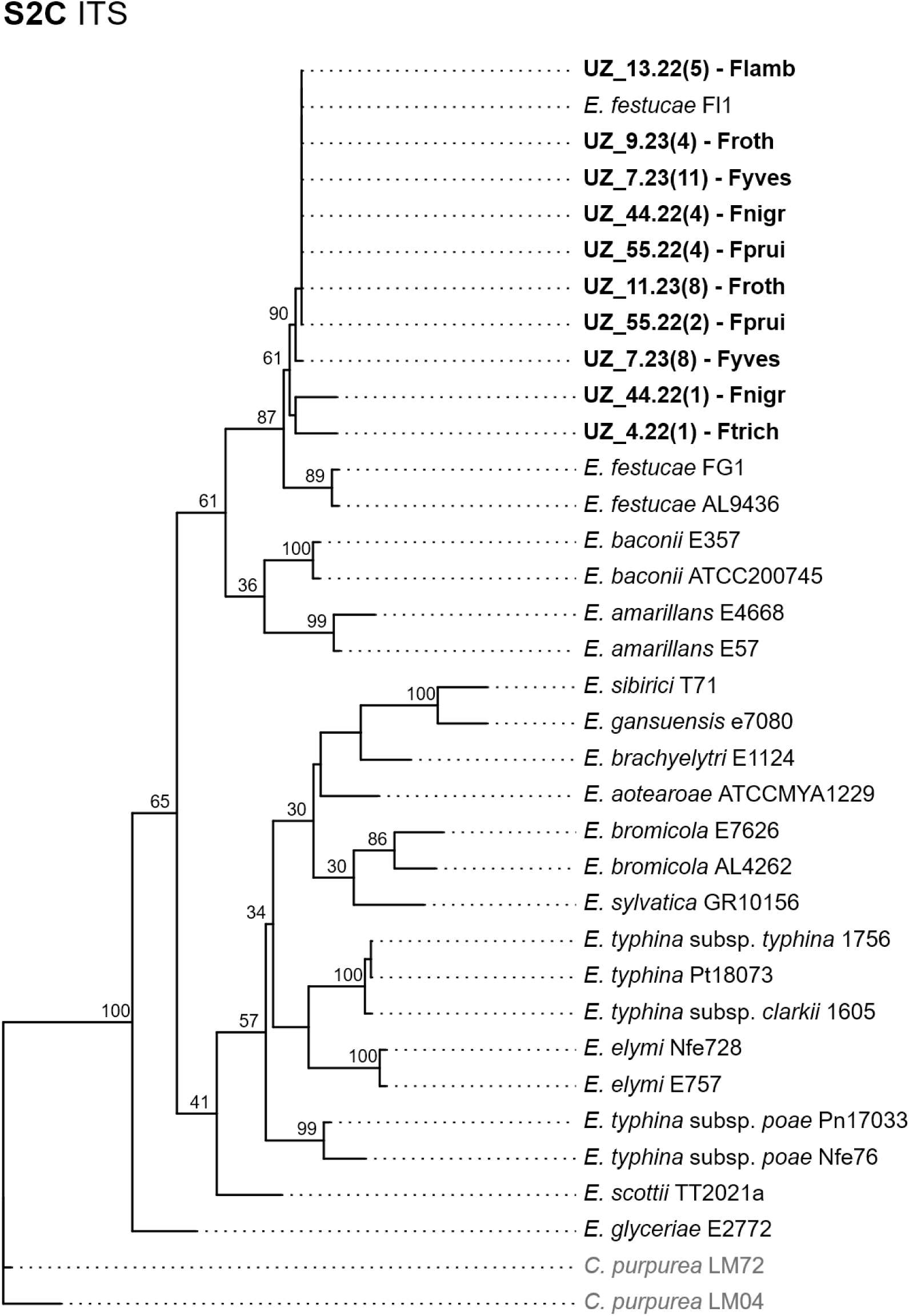
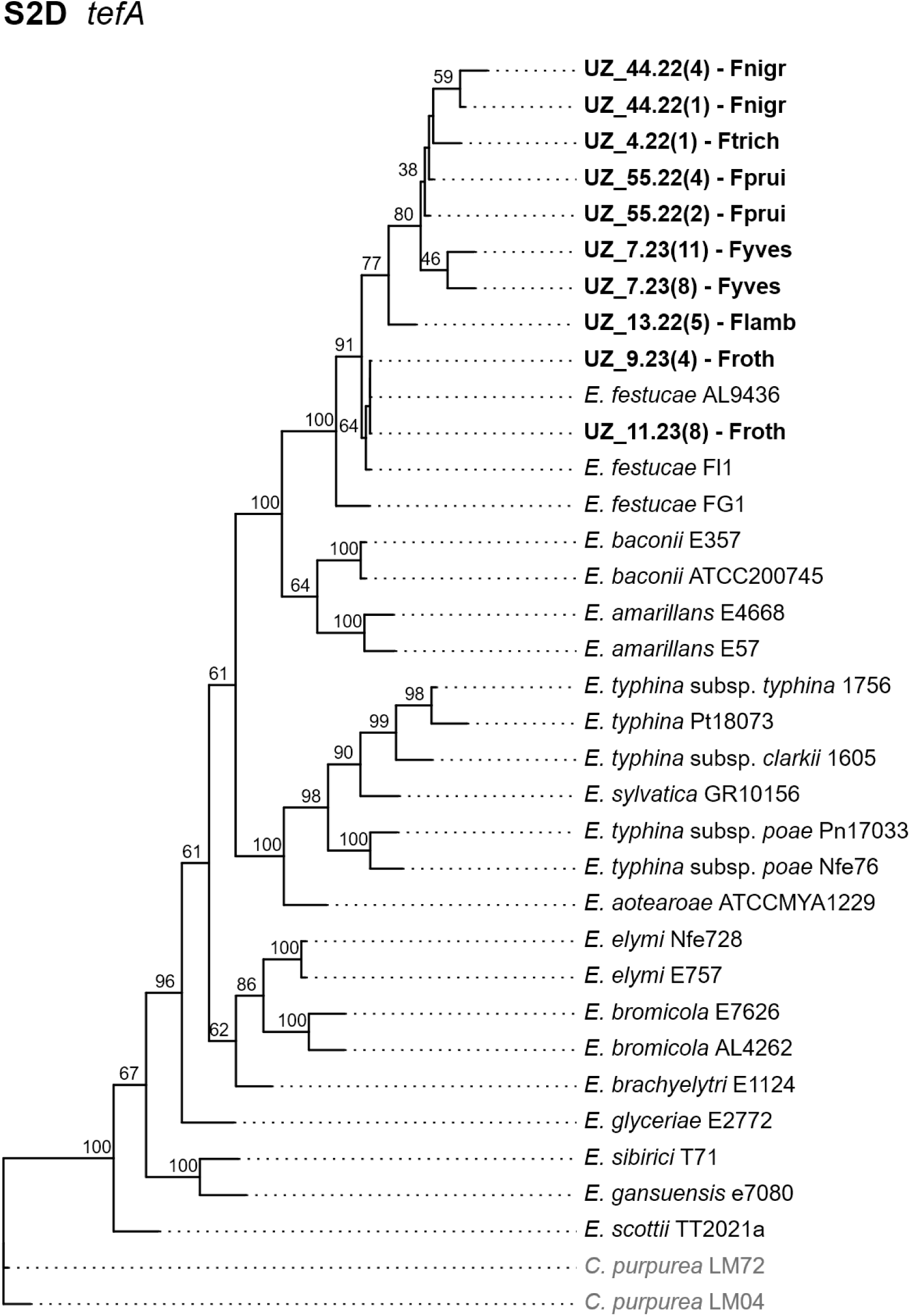
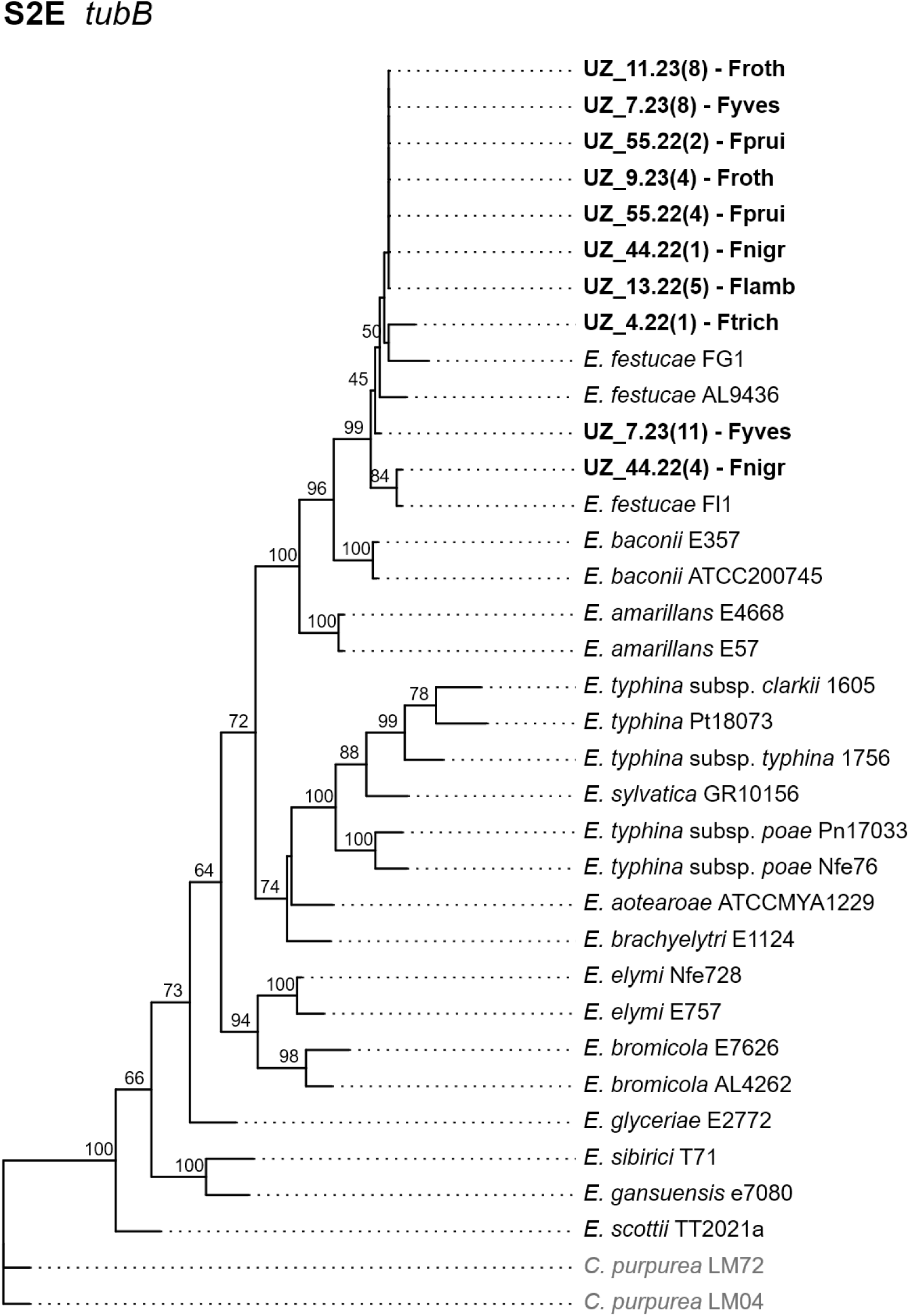
Maximum-likelihood phylogenetic trees (IQ-TREE2) of the studied *Epichloë festucae* strains based on five barcoding loci. **(A)** γ-actin (*actG*), **(B)** calmodulin (*CalM*), **(C)** rDNA ITS region, **(D)** translation elongation factor 1-α (*tefA*), and **(E)** β-tubulin (*tubB*). Analyzed samples are highlighted in bold (Table S9), and external sequences of other *Epichloë* specimens and the outgroup *Claviceps purpurea* were obtained from NCBI (Sotomayor-Alge et al., 2025). Node values represent ultrafast bootstrap support.

**Figure S3.**
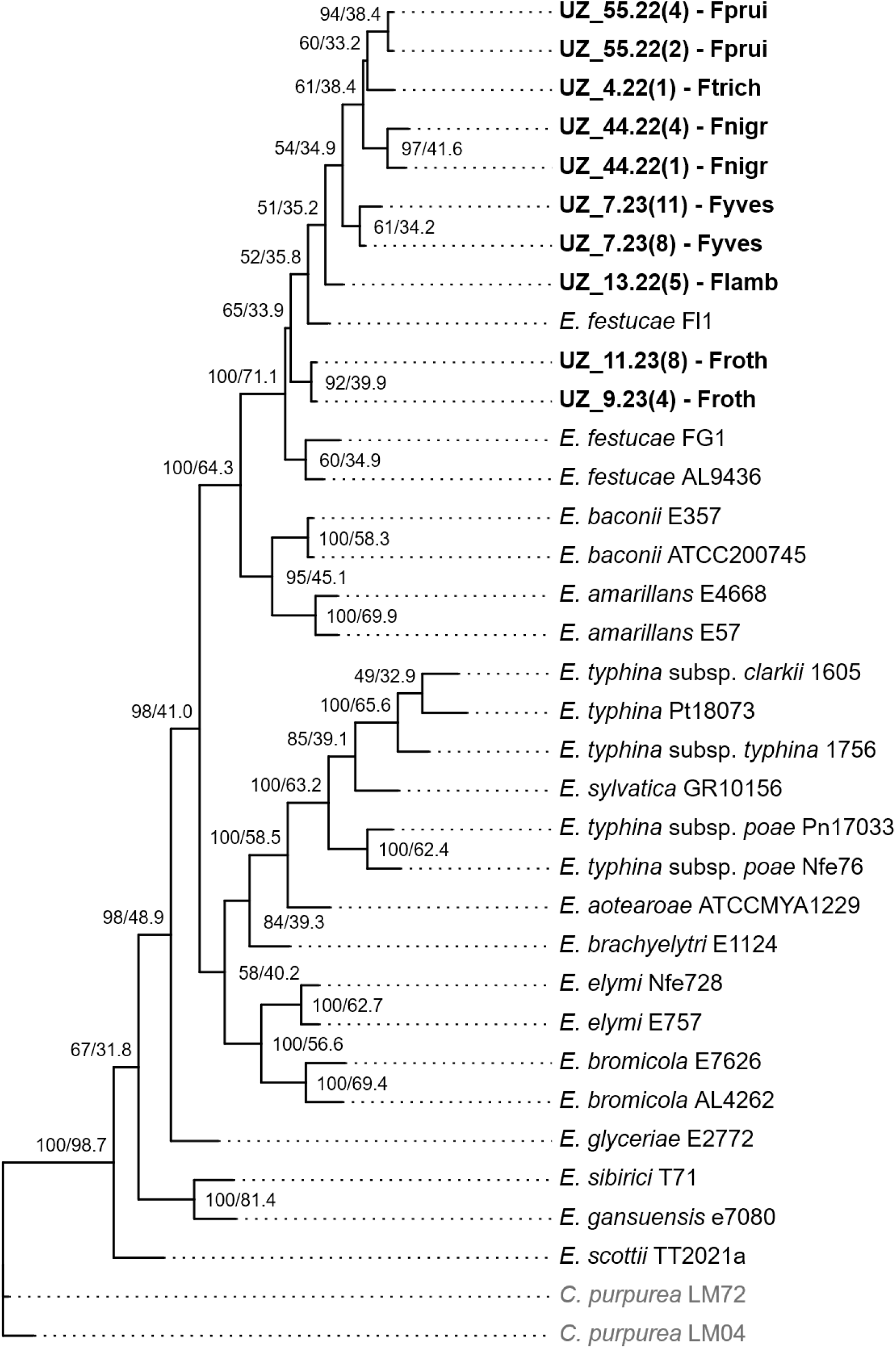
Concatenated maximum-likelihood phylogeny (IQ-TREE2) of the studied *Epichloë festucae* strains based on five nuclear loci (*actG*, *CalM*, ITS region, *tefA*, and *tubB*). Analyzed samples are highlighted in bold (Table S9) and external sequences of other *Epichloë* specimens and the outgroup *Claviceps purpurea* were obtained from NCBI (Sotomayor-Alge et al., 2025). Branch labels correspond to ultrafast bootstrap support (BS) / likelihood-based site concordance factor (sCFL).

## Supplementary tables

**Table S1.**
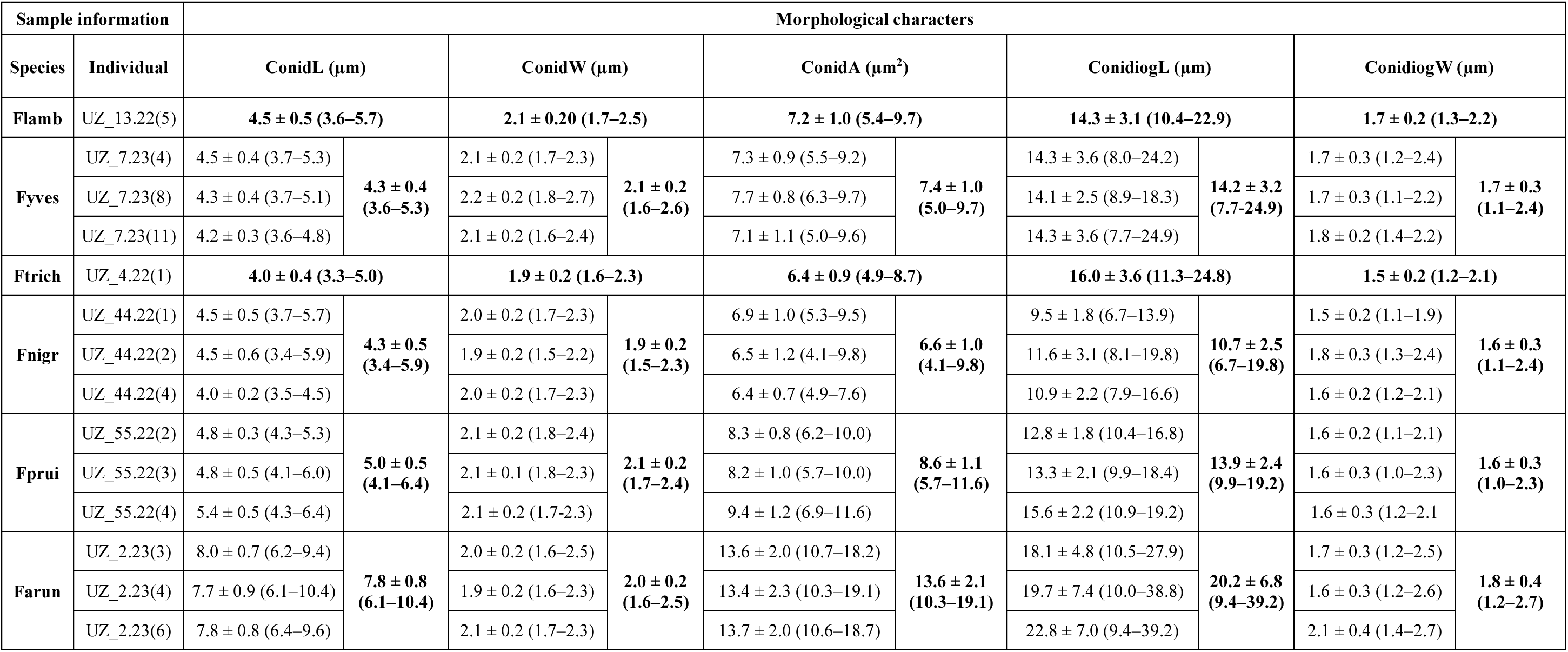
Morphological data of asexual characters (conidia and conidiogenous cells) of *Epichloë festucae* isolates from five fine-leaved *Festuca* host plants and of *Epichloë coenophiala* from one broad-leaved *Festuca* host plant. Holobionts analyzed belong to *Festuca* sect. *Festuca* [*F. lambinonii* (Flamb) and *F. yvesii* (Fyves)], *Festuca* sect. *Aulaxyper* [*Festuca trichophylla* (Ftrich), *Festuca nigrescens* (Fnigr), and *Festuca rubra* subsp. *pruinosa* (Fprui)], and *Festuca* subgen. *Schedonorus* [*Festuca arundinacea* subsp. *arundinacea* (Farun)]. Measurements include conidial length (ConidL), conidial width (ConidW), conidial area (ConidA), length of conidiogenous cell (ConidiogL) and width of the conidiogenous basal cell (ConidiogW). Values correspond to mean ± SD at isolate (individual) and host species levels (minimum value**–**maximum value) whenever possible. See also Fig. 1A.

| Sample information |  | Morphological characters |  |  |  |  |  |  |  |  |  |
| --- | --- | --- | --- | --- | --- | --- | --- | --- | --- | --- | --- |
| Species | Individual | ConidL (μm) |  | ConidW (μm) |  | ConidA (μm<sup>2</sup>) |  | ConidiogL (μm) |  | ConidiogW (μm) |  |
| Flamb | UZ_13.22(5) | 4.5 ± 0.5 (3.6–5.7) |  | 2.1 ± 0.20 (1.7–2.5) |  | 7.2 ± 1.0 (5.4–9.7) |  | 14.3 ± 3.1 (10.4–22.9) |  | 1.7 ± 0.2 (1.3–2.2) |  |
| Fyves | UZ_7.23(4) | 4.5 ± 0.4 (3.7–5.3) | 4.3 ± 0.4 (3.6–5.3) | 2.1 ± 0.2 (1.7–2.3) | 2.1 ± 0.2 (1.6–2.6) | 7.3 ± 0.9 (5.5–9.2) | 7.4 ± 1.0 (5.0–9.7) | 14.3 ± 3.6 (8.0–24.2) | 14.2 ± 3.2 (7.7–24.9) | 1.7 ± 0.3 (1.2–2.4) | 1.7 ± 0.3 (1.1–2.4) |
|  | UZ_7.23(8) | 4.3 ± 0.4 (3.7–5.1) |  | 2.2 ± 0.2 (1.8–2.7) |  | 7.7 ± 0.8 (6.3–9.7) |  | 14.1 ± 2.5 (8.9–18.3) |  | 1.7 ± 0.3 (1.1–2.2) |  |
|  | UZ_7.23(11) | 4.2 ± 0.3 (3.6–4.8) |  | 2.1 ± 0.2 (1.6–2.4) |  | 7.1 ± 1.1 (5.0–9.6) |  | 14.3 ± 3.6 (7.7–24.9) |  | 1.8 ± 0.2 (1.4–2.2) |  |
| Ftrich | UZ_4.22(1) | 4.0 ± 0.4 (3.3–5.0) |  | 1.9 ± 0.2 (1.6–2.3) |  | 6.4 ± 0.9 (4.9–8.7) |  | 16.0 ± 3.6 (11.3–24.8) |  | 1.5 ± 0.2 (1.2–2.1) |  |
| Fnigr | UZ_44.22(1) | 4.5 ± 0.5 (3.7–5.7) | 4.3 ± 0.5 (3.4–5.9) | 2.0 ± 0.2 (1.7–2.3) | 1.9 ± 0.2 (1.5–2.3) | 6.9 ± 1.0 (5.3–9.5) | 6.6 ± 1.0 (4.1–9.8) | 9.5 ± 1.8 (6.7–13.9) | 10.7 ± 2.5 (6.7–19.8) | 1.5 ± 0.2 (1.1–1.9) | 1.6 ± 0.3 (1.1–2.4) |
|  | UZ_44.22(2) | 4.5 ± 0.6 (3.4–5.9) |  | 1.9 ± 0.2 (1.5–2.2) |  | 6.5 ± 1.2 (4.1–9.8) |  | 11.6 ± 3.1 (8.1–19.8) |  | 1.8 ± 0.3 (1.3–2.4) |  |
|  | UZ_44.22(4) | 4.0 ± 0.2 (3.5–4.5) |  | 2.0 ± 0.2 (1.7–2.3) |  | 6.4 ± 0.7 (4.9–7.6) |  | 10.9 ± 2.2 (7.9–16.6) |  | 1.6 ± 0.2 (1.2–2.1) |  |
| Fprui | UZ_55.22(2) | 4.8 ± 0.3 (4.3–5.3) | 5.0 ± 0.5 (4.1–6.4) | 2.1 ± 0.2 (1.8–2.4) | 2.1 ± 0.2 (1.7–2.4) | 8.3 ± 0.8 (6.2–10.0) | 8.6 ± 1.1 (5.7–11.6) | 12.8 ± 1.8 (10.4–16.8) | 13.9 ± 2.4 (9.9–19.2) | 1.6 ± 0.2 (1.1–2.1) | 1.6 ± 0.3 (1.0–2.3) |
|  | UZ_55.22(3) | 4.8 ± 0.5 (4.1–6.0) |  | 2.1 ± 0.1 (1.8–2.3) |  | 8.2 ± 1.0 (5.7–10.0) |  | 13.3 ± 2.1 (9.9–18.4) |  | 1.6 ± 0.3 (1.0–2.3) |  |
|  | UZ_55.22(4) | 5.4 ± 0.5 (4.3–6.4) |  | 2.1 ± 0.2 (1.7–2.3) |  | 9.4 ± 1.2 (6.9–11.6) |  | 15.6 ± 2.2 (10.9–19.2) |  | 1.6 ± 0.3 (1.2–2.1) |  |
| Farun | UZ_2.23(3) | 8.0 ± 0.7 (6.2–9.4) | 7.8 ± 0.8 (6.1–10.4) | 2.0 ± 0.2 (1.6–2.5) | 2.0 ± 0.2 (1.6–2.5) | 13.6 ± 2.0 (10.7–18.2) | 13.6 ± 2.1 (10.3–19.1) | 18.1 ± 4.8 (10.5–27.9) | 20.2 ± 6.8 (9.4–39.2) | 1.7 ± 0.3 (1.2–2.5) | 1.8 ± 0.4 (1.2–2.7) |
|  | UZ_2.23(4) | 7.7 ± 0.9 (6.1–10.4) |  | 1.9 ± 0.2 (1.6–2.3) |  | 13.4 ± 2.3 (10.3–19.1) |  | 19.7 ± 7.4 (10.0–38.8) |  | 1.6 ± 0.3 (1.2–2.6) |  |
|  | UZ_2.23(6) | 7.8 ± 0.8 (6.4–9.6) |  | 2.1 ± 0.2 (1.7–2.3) |  | 13.7 ± 2.0 (10.6–18.7) |  | 22.8 ± 7.0 (9.4–39.2) |  | 2.1 ± 0.4 (1.4–2.7) |  |

**Table S2.**
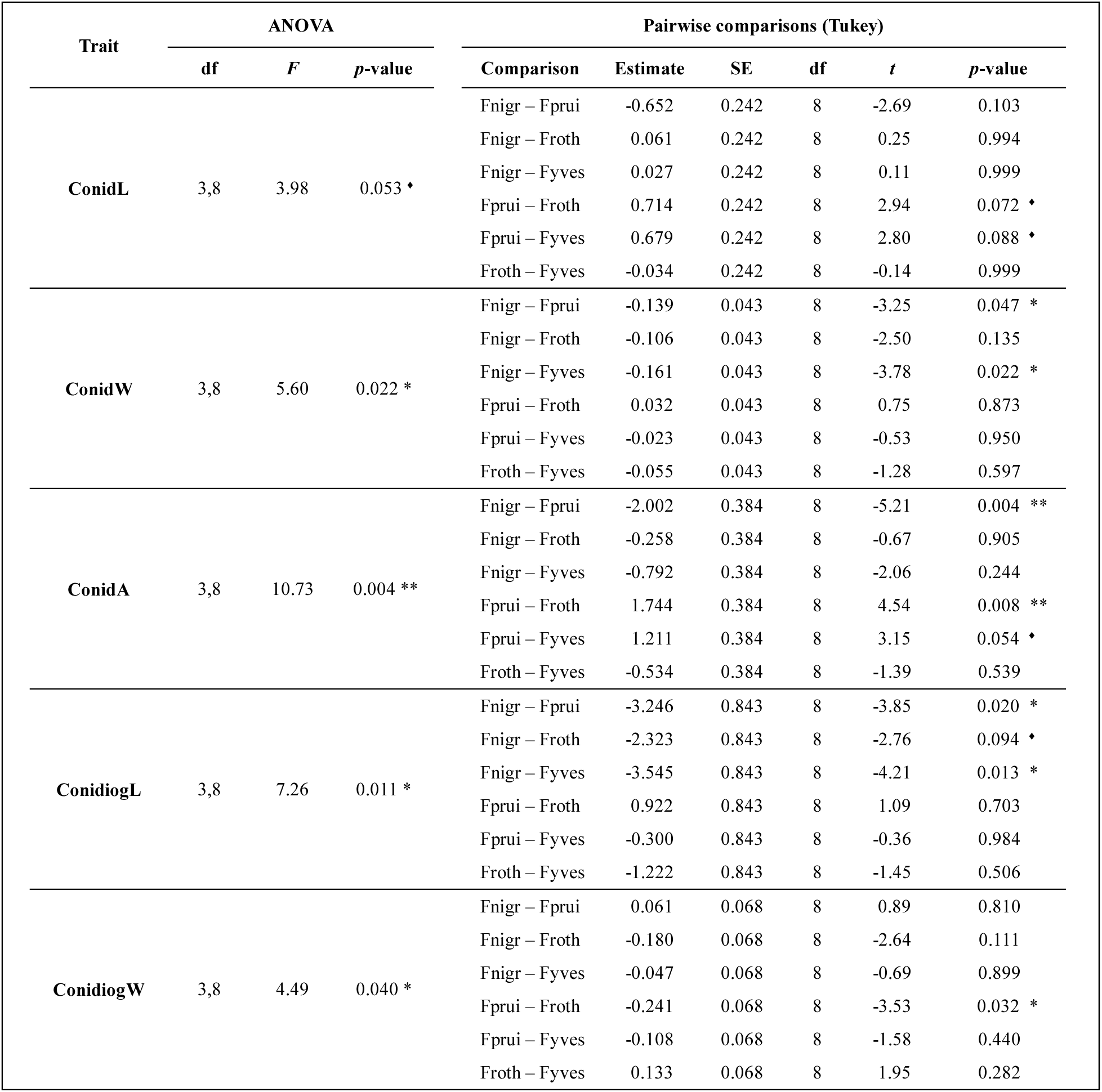
Results of linear mixed models (LMMs) testing the differences among *Epichloë festucae* strains in asexual characters (conidia and conidiogenous cells). Four host species were included in the analysis: *Festuca yvesii* (Fyves), *F. rothmaleri* (Froth), *F. nigrescens* (Fnigr) and *F. rubra* subsp. *pruinosa* (Fprui). Host species effect was evaluated using Type III ANOVA with Kenward–Roger approximation of degrees of freedom, and pairwise comparisons were performed using Tukey-adjusted estimated marginal means. For conidial width (ConidW), conidial area (ConidA), and conidiogenous cell length (ConidiogL), the model included isolate replicates nested within isolates (*Trait ∼ Host species + (1 | Ind/Rep)*), whereas for conidial length (ConidL) and conidiogenous cell width (ConidiogW) the model included a random intercept for isolate identity (*Trait ∼ Host species + (1 | Ind)*). Degrees of freedom were estimated using the Kenward–Roger approximation in ANOVA. Estimates represent pairwise differences between the means of *E. festucae* strains, along with their standard error (SE). Significance levels: 0.05 ≤ *p* ≤ 0.01 (*); 0.01 ≤ *p* ≤ 0.001 (**); *p* < 0.001 (***). The diamond indicates a marginally significant value.

**Table S3.** Variance partitioning and variance explained by linear mixed models (LMMs) fitted to each morphological trait of the *Epichloë festucae* strains. For conidial width (ConidW), conidial area (ConidA), and conidiogenous cell length (ConidiogL), the model included isolate replicates nested within isolate identity (*Trait ∼ Host species +* (*1 | Ind/Rep*)), whereas for conidial length (ConidL) and conidiogenous cell width (ConidiogW) it included a random intercept for isolate identity (*Trait ∼ Host species +* (*1 | Ind*)). Conditional (*R*²c) and marginal (*R*²m) coefficients of determination represent the variance explained by the full model (fixed and random effects) and by fixed effects alone, respectively. Values in parentheses next to R²m indicate the percentage of the explained variance attributable to fixed effects (i.e., (*R*²m / *R*²c) × 100). Variance components show the contribution of each random effect and residual variance to the total model variance. “Ind” denotes isolate (individual), “Rep:Ind” indicates isolate replicate nested within isolate and “Residual” represents the remaining unexplained variance not accounted for by the model.

| Trait | Variance explained ($R^2$) | | Variance components | | |
| --- | --- | --- | --- | --- | --- |
| | $R^2c$ | $R^2m$ (%) | Random effect | Variance | Proportion |
| ConidL | 0.51 | 0.27 (52.9 %) | Ind | 0.083 | 0.34 |
|  |  |  | Residual | 0.161 | 0.66 |
| ConidW | 0.26 | 0.12 (46.2 %) | Ind | 0.001 | 0.03 |
|  |  |  | Rep:Ind | 0.033 | 0.12 |
|  |  |  | Residual | 0.023 | 0.85 |
| ConidA | 0.48 | 0.35 (72.9 %) | Ind | 0.177 | 0.16 |
|  |  |  | Rep:Ind | 0.045 | 0.04 |
|  |  |  | Residual | 0.885 | 0.80 |
| ConidiogL | 0.38 | 0.21 (55.3 %) | Ind | 0.536 | 0.08 |
|  |  |  | Rep:Ind | 1.025 | 0.14 |
|  |  |  | Residual | 5.646 | 0.78 |
| ConidiogW | 0.13 | 0.09 (69.2 %) | Ind | 0.004 | 0.05 |
|  |  |  | Residual | 0.081 | 0.95 |

**Table S4.** Multivariate statistical analysis of morphometric variation among *Epichloë festucae* strains associated with different *Festuca* host species. **(A)** Global PERMANOVA testing the effect of host species on standardized conidial morphometric variables of their *Epichloë festucae* strains at three hierarchical levels (spores, plates and isolates). Four host species were included in the analysis: *Festuca yvesii* (Fyves), *F. rothmaleri* (Froth), *F. nigrescens* (Fnigr) and *F. rubra* subsp. *pruinosa* (Fprui). Distances were calculated using Euclidean distance on standardized variables with 10,000 permutations. PERMDISP tests were performed to assess homogeneity of multivariate dispersions among host species. **(B)** Pairwise PERMANOVA comparisons among host species based on Euclidean distances of the same standardized morphometric variables of their *Epichloë festucae* strains at the isolate level. Each comparison involved three isolates per host species (n = 3 per species). *P*-values were adjusted for multiple comparisons using the Holm method. PERMDISP tests were conducted for each pairwise comparison to evaluate differences in multivariate dispersion. Due to the limited number of isolates per species, the number of possible permutations was restricted to 20, resulting in fixed *p*-values across pairwise comparisons. Significance level: *p* < 0.05 (*).

| (A) | Predictor | GLOBAL PERMANOVA |  |  |  |  | PERMDISP test |  |
| --- | --- | --- | --- | --- | --- | --- | --- | --- |
| | | N | df residual | $R^2$ | pseudo-$F$ | $p$-value | $F$ | $p$-value |
|  | All measurements (exploratory) | 360 | 356 | 0.18 | 26.23 | 1.000 | 0.90 | 0.448 |
|  | Mean per plate (exploratory) | 36 | 32 | 0.43 | 8.19 | 1.000 | 1.79 | 0.170 |
|  | Mean per isolate (main result) | 12 | 8 | 0.66 | 5.15 | < 0.001 * | 0.51 | 0.707 |

| (B) | Comparison | PAIRWISE PERMANOVA |  |  |  |  | PERMDISP test |  |
| --- | --- | --- | --- | --- | --- | --- | --- | --- |
| | | N | $R^2$ | pseudo-$F$ | $p$-value | $p$-adj | $F$ | $p$-value |
|  | Fnigr – Fprui | 3 vs 3 | 0.59 | 5.87 | 0.1 | 0.6 | 0.66 | 0.401 |
|  | Fnigr – Froth | 3 vs 3 | 0.51 | 4.21 | 0.1 | 0.6 | 2.19 | 0.301 |
|  | Fnigr – Fyves | 3 vs 3 | 0.58 | 5.54 | 0.1 | 0.6 | 1.28 | 0.301 |
|  | Fprui – Froth | 3 vs 3 | 0.68 | 8.56 | 0.1 | 0.6 | 0.00 | 0.901 |
|  | Fprui – Fyves | 3 vs 3 | 0.48 | 3.67 | 0.1 | 0.6 | 0.04 | 0.801 |
|  | Froth – Fyves | 3 vs 3 | 0.41 | 2.73 | 0.1 | 0.6 | 0.20 | 0.401 |

**Table S5.**
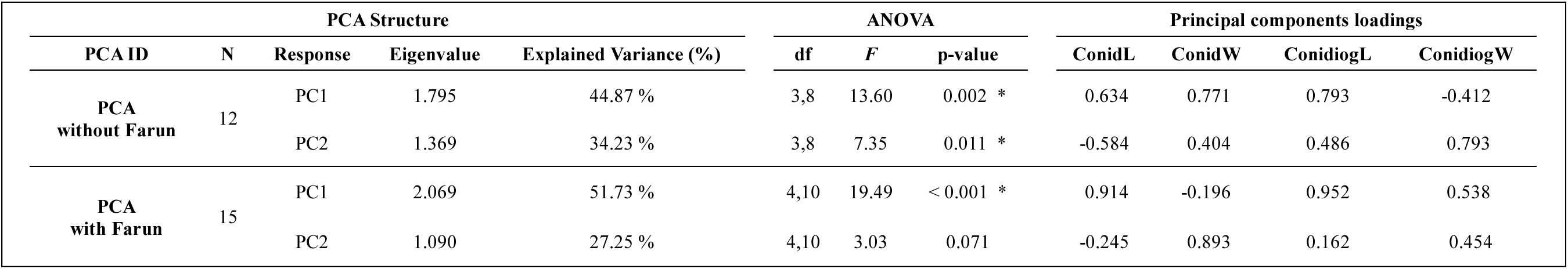
Statistical results associated with the Principal Component Analysis of conidial and conidiogenous cell morphometric traits in *Epichloë festucae*. Values correspond to mean measurements per isolate from holobionts of *Festuca yvesii* (Fyves), *F. rothmaleri* (Froth), *F. nigrescens* (Fnigr) and *F. rubra* subsp. *pruinosa* (Fprui) excluding (n = 12) and including (n = 15) samples from the holobiont of *F. arundinacea* subsp. *arundinacea* – *E. coenophiala* (Farun). For each PCA, eigenvalues, percentage of explained variance, ANOVA results testing for differences among species along the first two principal components (PC1 and PC2) and principal components loadings are reported. Loadings represent correlations between the morphometric variables –conidial length (ConidL), conidial width (ConidW), conidiogenous cell length (ConidiogL), and conidiogenous cell width (ConidiogW)– and the corresponding principal component axes. Significance level: *p* < 0.05 (*). See also Fig. 1B.

**Table S6.**
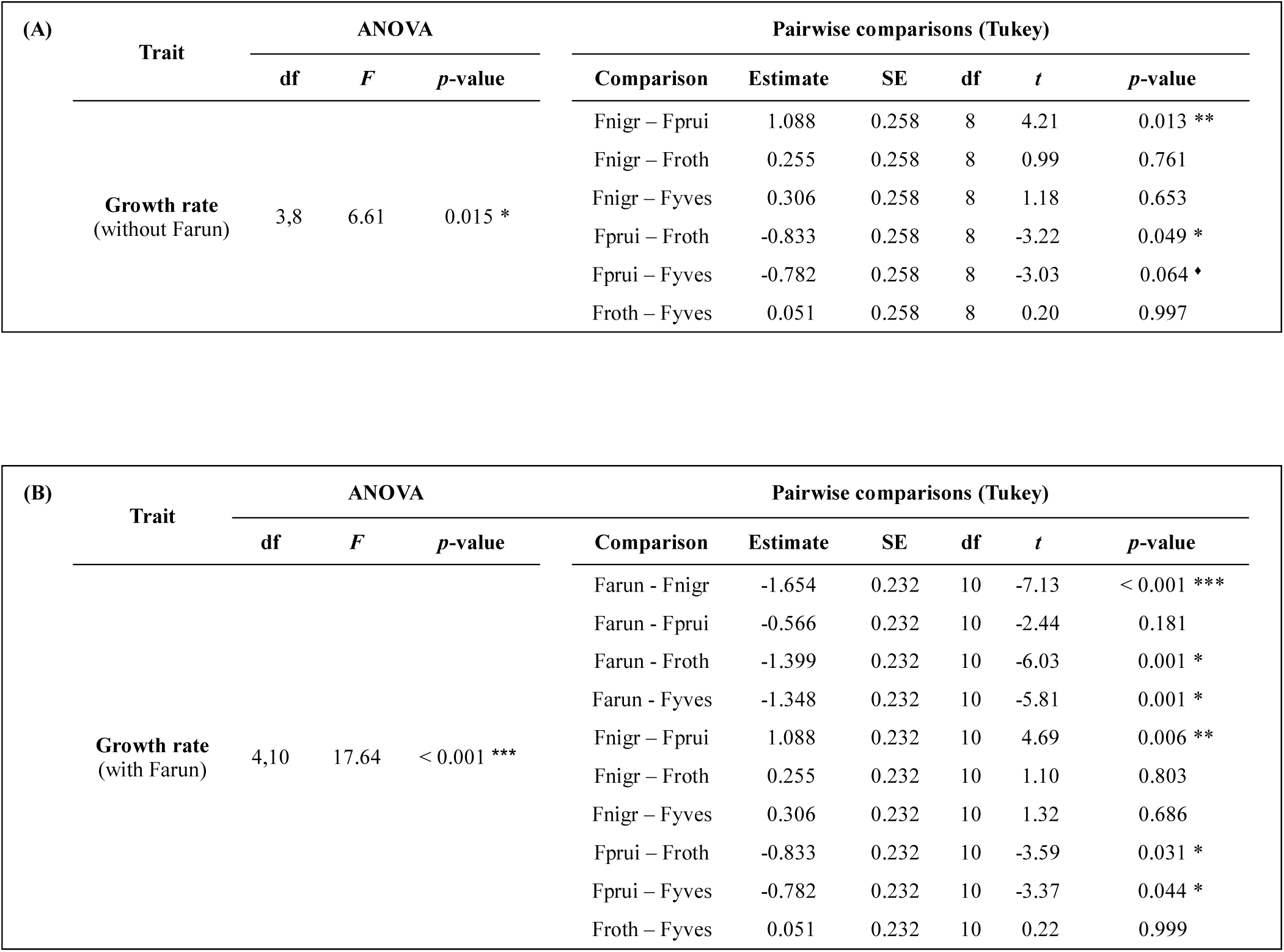
Linear mixed-effects models of growth rate variation (mm/day) among *Epichloë* isolates. Results among *Epichloë festucae* isolates associated with the holobionts Fnigr, Fprui, Fyves, and Froth **(A)**, and including the species *Epichloë coenophiala* isolated from the holobiont Farun **(B)**. Each host species was represented by three *Epichloë* isolates. Host species was treated as a fixed effect, with isolate identity nested within host species included as a random effect following the model structure *GR ∼ Host species + (1 | Host species:Ind)*. Degrees of freedom for the ANOVA were estimated using the Kenward–Roger approximation. Pairwise differences among species were assessed using Tukey-adjusted contrasts based on estimated marginal means. Estimates represent pairwise differences between strain means with their associated standard errors (SE). Significance levels: 0.05 ≤ *p* ≤ 0.01 (*); 0.01 ≤ *p* ≤ 0.001 (**); *p* < 0.001 (***). The diamond indicates a marginally significant value.

**Table S7.** Quartet support values for the phylogenetic ASTRAL coalescent tree of *Epichloë festucae* strains. Values of q1 (main topology), q2 (first alternative topology), and q3 (second alternative topology) are reported for each node of the species tree inferred using ASTRAL-III. See also Fig. 3.

| Node | q1 | q2 | q3 |
| --- | --- | --- | --- |
| 1 | 1 | 0 | 0 |
| 2 | 0.76 | 0.11 | 0.13 |
| 3 | 1 | 0 | 0 |
| 4 | 0.64 | 0.32 | 0.04 |
| 5 | 0.76 | 0.04 | 0.2 |
| 6 | 0.67 | 0.26 | 0.07 |
| 7 | 0.47 | 0.24 | 0.29 |
| 8 | 0.85 | 0.09 | 0.05 |
| 9 | 1 | 0 | 0 |
| 10 | 1 | 0 | 0 |
| 11 | 0.81 | 0.1 | 0.08 |
| 12 | 0.83 | 0.15 | 0.01 |
| 13 | 1 | 0 | 0 |
| 14 | 0.54 | 0.24 | 0.21 |
| 15 | 1 | 0 | 0 |
| 16 | 0.4 | 0.4 | 0.2 |
| 17 | 1 | 0 | 0 |
| 18 | 0.8 | 0 | 0.2 |
| 19 | 1 | 0 | 0 |
| 20 | 1 | 0 | 0 |
| 21 | 1 | 0 | 0 |
| 22 | 0.65 | 0.06 | 0.29 |
| 23 | 0.43 | 0.2 | 0.37 |
| 24 | 0.7 | 0.28 | 0.02 |
| 25 | 0.5 | 0.18 | 0.33 |
| 26 | 0.58 | 0.04 | 0.38 |
| 27 | 0.46 | 0.41 | 0.14 |
| 28 | 0.39 | 0.29 | 0.32 |
| 29 | 0.56 | 0.21 | 0.23 |
| 30 | 0.47 | 0.24 | 0.29 |
| 31 | 0.73 | 0.25 | 0.01 |
| 32 | 0.41 | 0.21 | 0.38 |

**Table S8.**
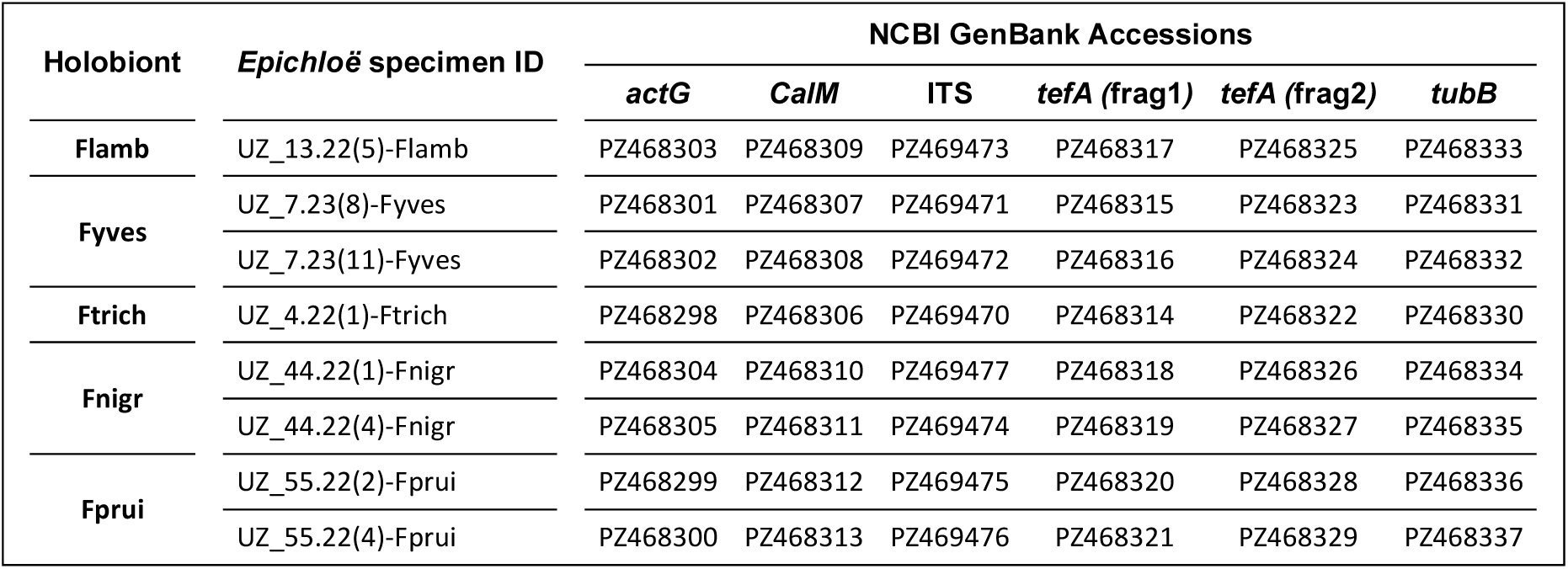
GenBank accession numbers of sequenced *Epichloë festucae* loci. Five barcoding nuclear loci *actG*, *CalM*, ITS, *tefA* (frag1 and frag2) and *tubB* were sequenced from the *Epichloë festucae* strains under study isolated from their respective fine-leaved *Festuca* host plants.

**Table S9.**
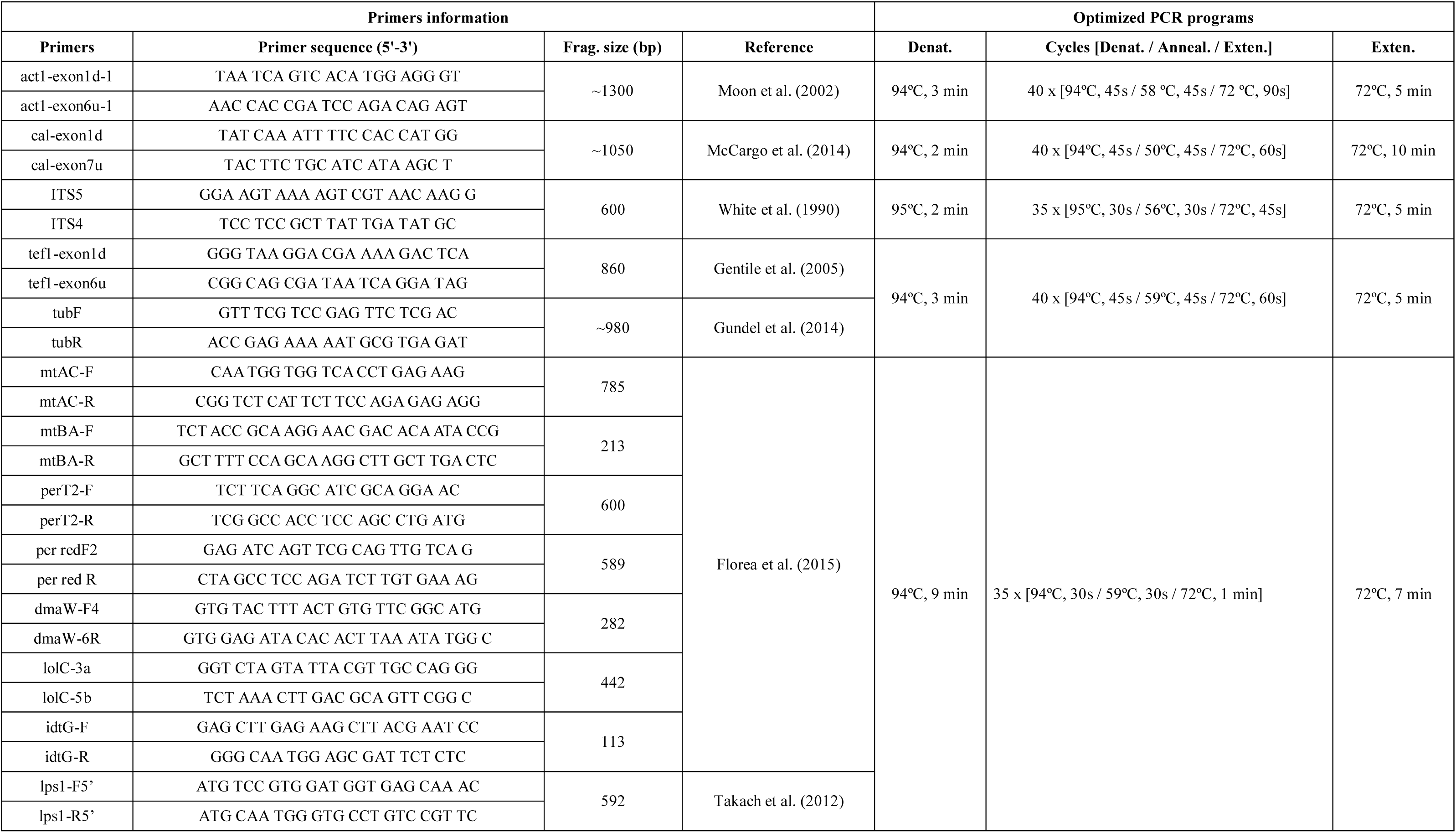
Primer pairs and PCR amplification conditions used in this study. Amplifications conducted on five barcoding loci [*actG*, *CalM*, ITS, *tef1* and *tub2*], endophyte mating types [*MTA*(*mtAC* and *MTB* (*mtBA*)] and main alkaloid synthetic pathways genes [pyrrolopyrazines (*perT2* and *per red*), ergot alkaloids (*dmaW* and *lpsA*), 1-aminopyrrolizidines (*lolC*), and indole-diterpenes (*idtG*)]. References stand for the origin of the primers and original PCR programs used. Optimized PCR programs follow the structure initial denaturation (Denat.), PCR cycles (Cycles [Denat. / Anneal. / Exten.]) and final extension (Exten.) within the table.

